# Obscurin Rho GEF domains are phosphorylated by MST-family kinases but do not exhibit nucleotide exchange factor activity towards Rho GTPases *in vitro*

**DOI:** 10.1101/2022.10.14.508903

**Authors:** Daniel Koch, Ay Lin Kho, Atsushi Fukuzawa, Alexander Alexandrovich, Andrew Beavil, Mark Pfuhl, Martin Rees, Mathias Gautel

## Abstract

Obscurin is a giant muscle protein (>800 kDa) featuring multiple signalling domains, including an SH3-DH-PH domain triplet from the Trio-subfamily of guanosine nucleotide exchange factors (GEFs). While previous research suggests that these domains can activate the small GTPases RhoA and RhoQ in cells, *in vitro* characterization of these interactions using biophysical techniques has been hampered by the intrinsic instability of obscurin GEF domains. To study substrate specificity, mechanism and regulation of obscurin GEF function by individual domains, we successfully optimized recombinant production of obscurin GEF domains and found that MST-family kinases phosphorylate the obscurin DH domain at Thr5798. Despite extensive testing of multiple GEF domain fragments, we did not detect any nucleotide exchange activity *in vitro* against 9 representative small GTPases. Bioinformatic analyses show that obscurin differs from other Trio-subfamily GEFs in several important aspects. While further research is necessary to evaluate obscurin GEF activity *in vivo*, our results indicate that obscurin has atypical GEF domains that, if catalytically active at all, are subject to complex regulation.

## 1. Introduction

Obscurin is a giant muscle protein of yet poorly understood functions ^1^. Its N-terminus links the proteins titin and myomesin at the sarcomeric M-band and the C-terminus of obscurin isoform A binds to small ankyrins at the sarcoplasmic reticulum membrane and at subsarcolemmal protein complexes, thereby creating multiple physical links that contribute to M-band stability and membrane integrity ^2–8^. In addition to these structural roles, obscurin is likely involved in multiple signalling contexts as it features two kinase domains, an IQ-motif, two calmodulin binding regions and an SH3-DH-PH RhoGEF domain triplet ^1,9^. Small GTPases of the Rho-family are a class of important signalling proteins that cycle between a biologically inactive GDP-bound form and a functional GTP-bound form and regulate diverse cellular functions including gene expression, cytoskeletal remodelling and contractility ^10^. Similar to phosphorylation being catalyzed by a kinase, transition to the active GTP state is promoted by guanosine nucleotide exchange factors (GEFs) ^11^. Previous studies showed that the obscurin DH-PH domains may act as a GEF on GTPases RhoA and RhoQ/TC10 but not Rac1 or Cdc42 and that these interactions are important for myofibril organization and may alter gene expression in response to mechanical signals ^12,13^. However, these insights were obtained using co-IP techniques involving nucleotide-state modifying Rho mutants and effector-domain pulldown assays from cell homogenates. While these approaches are suitable for a first characterization, they offer little information on substrate specificity and kinetics and can be prone to unintended consequences resulting from GTPase-mutants, overexpression of the GEF and unidentified cellular factors ^14,15^. Characterising the regulation and mechanism of GEF activity itself is subject to similar experimental constraints. Moreover, obscurin has only been tested against four GTPases although the Rho-family features more than 20 members, many of which could be potential substrates of the obscurin RhoGEF domains ^10,16^.

In contrast, biochemical, biophysical and structural *in vitro* methods allow for quantitative and precisely controlled characterization of kinetics, mechanism and regulation of GTPase:GEF interactions. Here, we report the production of soluble, stable recombinant obscurin GEF domains necessary for such studies. We identified kinases and phosphatases that can regulate the phosphorylation status of the obscurin RhoGEF region at a physiological site and tested over 10 different obscurin RhoGEF fragments against RhoA, RhoB, RhoC, Rac1, Rac2, Rac3, RhoG, Cdc42 and RhoQ/TC10 to determine kinetics, substrate specificity and regulation of the obscurin DH domain by adjacent domains and phosphorylation. Surprisingly, we found that none of the obscurin RhoGEF fragments exhibited any GEF activity. Bioinformatic analyses suggest that obscurin is an atypical member of the trio-subfamily of RhoGEFs to which it belongs and differs from other subfamily members at several important and conserved residues implicated in interactions with substrate GTPases, potentially explaining the lack of activity *in vitro*. We discuss implications for our understanding of obscurin RhoGEF function and outline avenues for future research.

## 2. Results

### Construct design and purification of human obscurin RhoGEF domains

Initial attempts to purify recombinant, His_6_-tagged obscurin RhoGEF domains (SH3-DH-PH, DH-PH, DH and PH) following expression in *E. coli* were not successful in our hands. Domain boundaries of these constructs were based on exon boundaries. We were able, however, to successfully express and purify obscurin SH3 and SH3-DH at high purity and at a yield of >5-10 mg/L culture (**Figure S1**). Since these protein fragments were stable at high concentrations without precipitation, we reasoned that the insolubility of the catalytic DH domain may be a result of suboptimal domain boundaries. We thus performed a bioinformatic analysis of the amino acid sequence of obscurin RhoGEF domains using multiple sequence and structure prediction algorithms and found that strict adherence to exon boundaries may disrupt the first N-terminal alpha-helix of the obscurin DH domain and excludes several highly conserved residues (**Figure 1A**). Using this information, we designed and screened 39 additional constructs of the DH, PH and DH-PH domains and found that DH domain constructs that fully include the predicted N-terminal extension resulted in soluble protein fragments at the correct molecular weight in small-scale purification assays **(Figure S2-3**). DH-PH or PH domain constructs, however, did not result in soluble protein, despite different domain boundaries, indicating that the PH domain is intrinsically unstable and difficult to purify. Larger scale purification of two selected DH domain fragments resulted in milligrams of highly pure, stable and folded protein (**Figure 1B-C and S4**).

**Figure 1.**
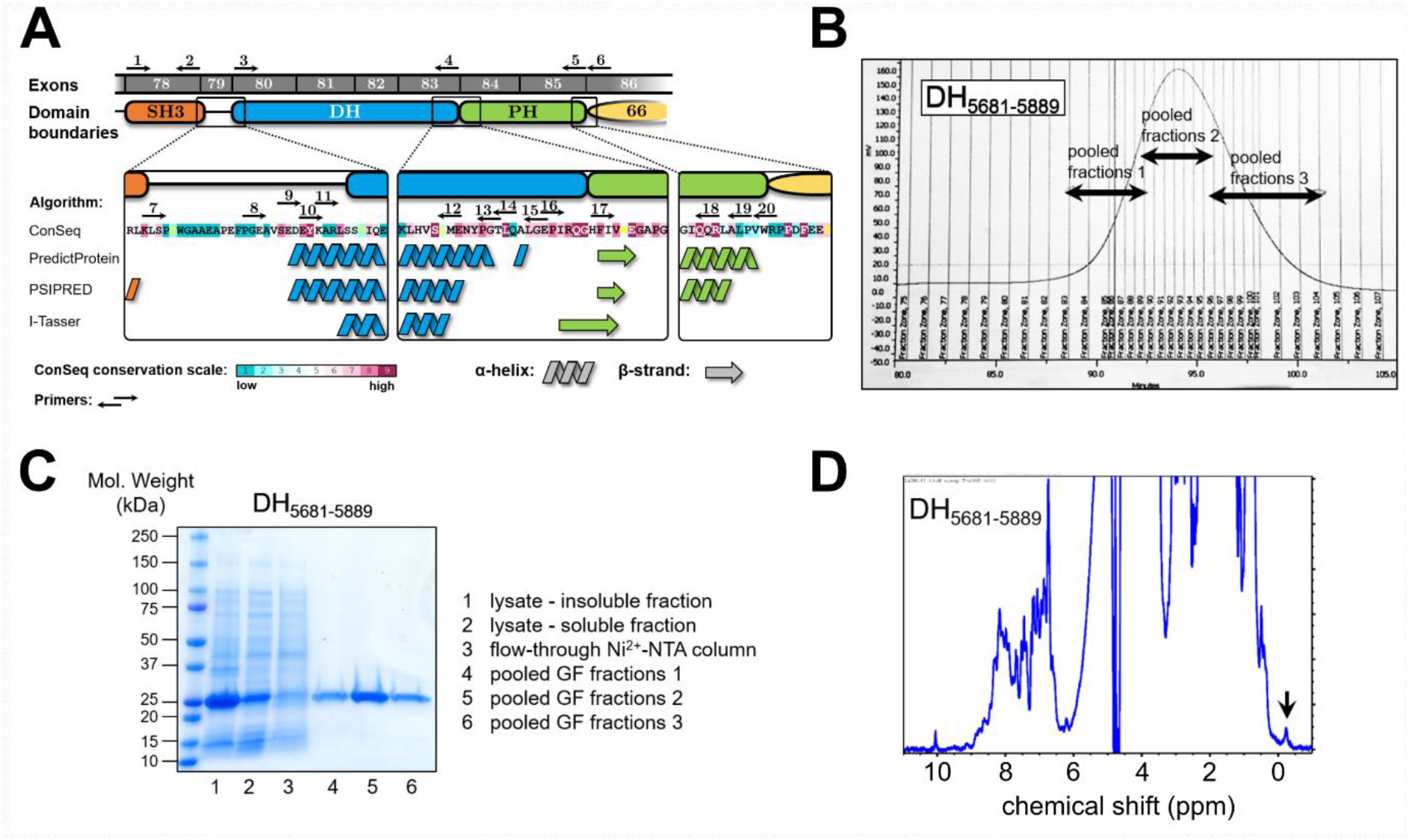
**A**, obscurin RhoGEF region. Exon boundaries and bioinformatic analysis and structure prediction based on amino acid sequence of human obscurin B isoform (NCBI ref. seq. NM_001098623). **B**, gel-filtration elution profile of DH domain fragment comprising residues 5681-5889. **C**, SDS-PAGE analysis of samples at different steps of the purification process of the DH_5681-5889_ fragment. **D**, 1D-NMR analysis of purified DH_5681-5889_ fragment. NMR spectrum exhibits wide peak dispersal and peaks below 0 ppm (black arrow), indicating that the protein is folded.

### Human obscurin RhoGEF domains do not exhibit catalytic activity towards Rho GTPases in vitro

The availability of highly pure and folded protein domains prompted us to characterize the catalytic activity of the obscurin RhoGEF domains towards Rho GTPases using enzyme kinetic methods. Using GTPases prepared with the fluorescent nucleotide 2’/3’-O-(N-Methyl-anthraniloyl)-GDP as a substrate, we tested whether obscurin SH3-DH and DH fragments can facilitate nucleotide exchange on any of the typical Rho GTPases that cycle between GDP/GTP states and are thus regulated by GEFs (with the exception of RhoJ, which was not soluble in our hands). Surprisingly, we found that the nucleotide exchange rate after addition of obscurin RhoGEF fragments was indistinguishable from the intrinsic nucleotide exchange rate – in stark contrast to the positive controls (**Figure 2** and **S5**). In other words, neither of the obscurin RhoGEF fragments exhibited nucleotide exchange activity towards any of the tested Rho GTPases, including RhoA and RhoQ/TC10.

**Figure 2.**
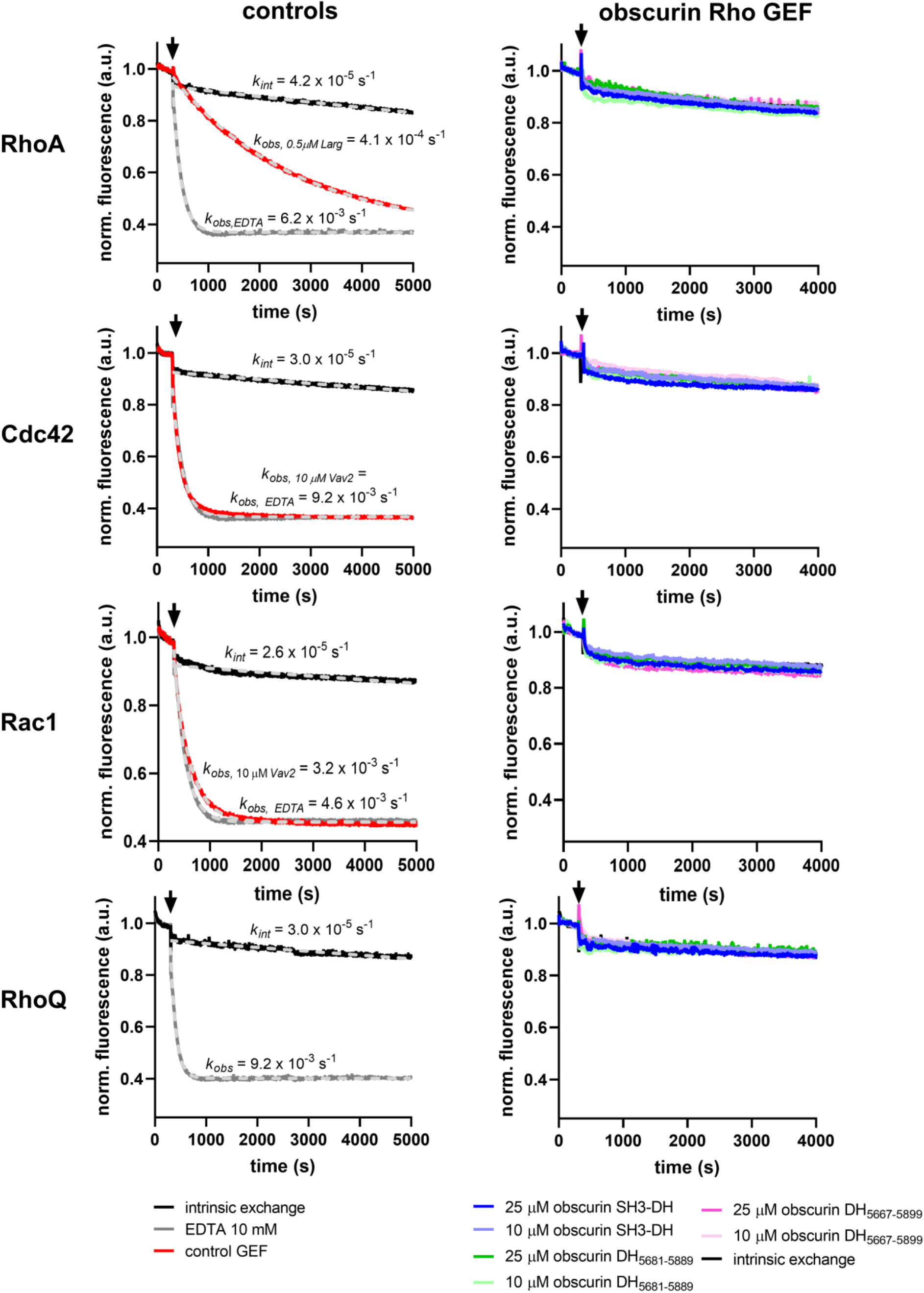
Guanosine nucleotide exchange factor activity of obscurin RhoGEF fragments towards RhoA, Cdc42, Rac1 and RhoQ/TC10. Left panels show the intrinsic nucleotide activity as a negative control, EDTA and GEFs Vav2 and Larg as positive controls for each GTPase. Right panels show the nucleotide exchange activity after addition of obscurin. Black arrows indicate addition of buffer/GEF/EDTA. Data represent mean of n = 2-3 experiments.

To confirm these results, we performed different pull-down experiments using a previously described protocol to stabilize the nucleotide-free high-affinity GEF-GTPase complex to specifically enrich GEFs ^17^. Using the putative substrate RhoA as a bait, we found that regardless of whether RhoA is bound to nucleotides, the bare DH domain does not bind to RhoA, suggesting RhoA is neither a substrate, nor binding partner of obscurin’s DH domain *in vitro* (**Figure S6**). Our findings thus disagree with previous reports of obscurin RhoGEF domains activating RhoA and RhoQ/TC10 ^12,13^.

To understand the reason for the discrepancy posed by our results, we further analyzed the sequence and predicted structure of obscurin RhoGEF domains, allowing us to better interpret our results in the context of the currently available mechanistic paradigms for Rho GEFs. The general mechanisms of DH-domain GEFs are well characterized and reviewed in the literature ^11,18–20^. Due to the amino acid sequence of its RhoGEF domains, obscurin belongs to the Trio-subfamily of RhoGEFs, which includes the GEFs dbl and dbs, for which detailed structural and kinetic data are available ^21^. Interestingly, the DH domains of dbl, dbs and trio have a much lower nucleotide exchange factor activity in the absence of their corresponding PH domain ^22^.

Subsequent work demonstrated that residues at the DH-PH domain interface (including residues from the PH domain) make contact with the switch II region of the substrate GTPase and contribute to the catalytic activity of the GEF (**Figure 3A, B**) ^19,20^. These residues are also present in most other trio-subfamily GEFs and are conserved in the obscurin sequence in humans, mice, chicken and zebrafish, but not in drosophila or nematodes (**Figure S7**) ^19^.

**Figure 3.**
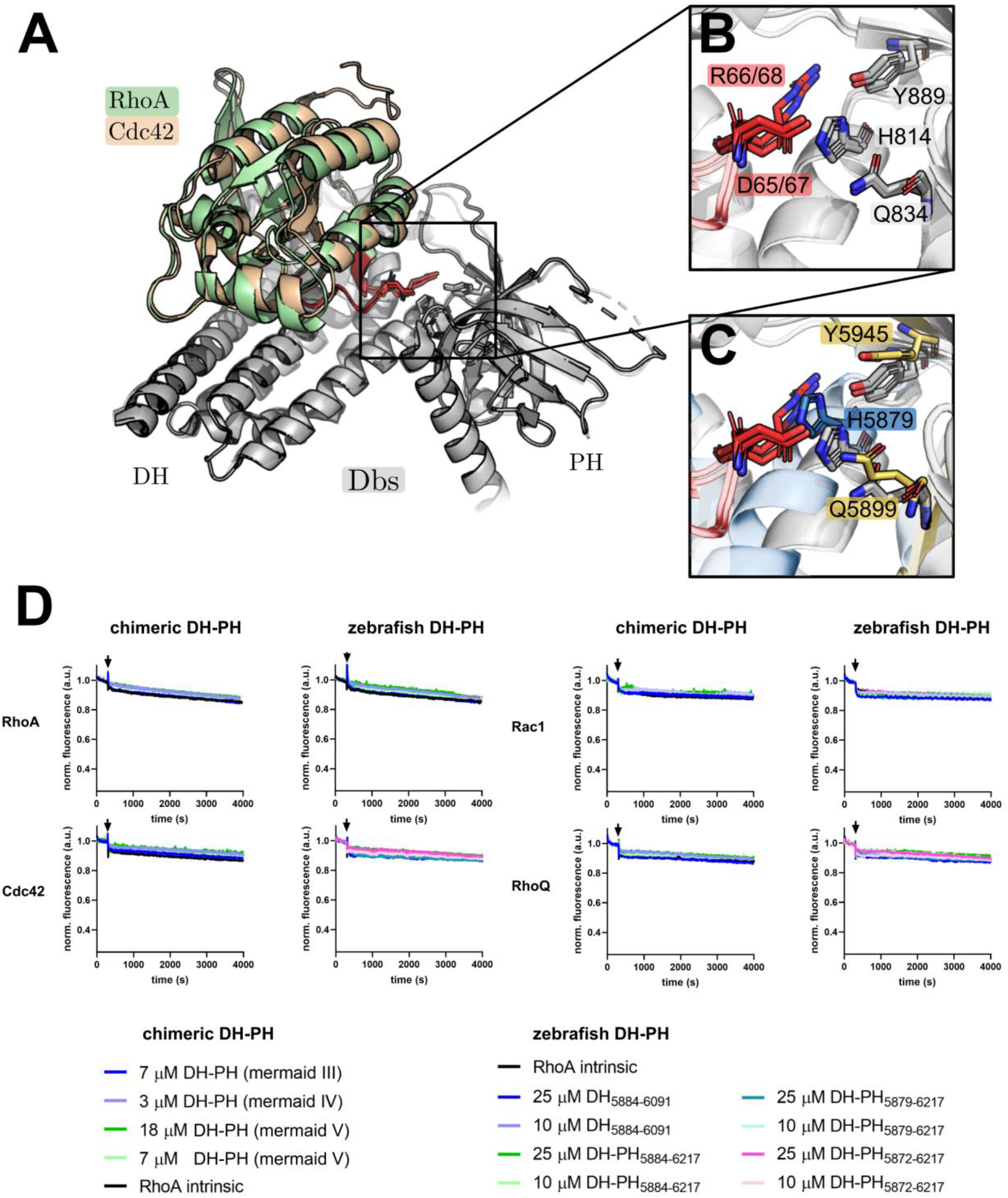
**A-C**, Structural determinants of nucleotide exchange activity in Trio-subfamily GEFs. **A**, the aligned structures of Dbs bound to RhoA (PDB-ID: 1LB1) and Cdc42 (PDB-ID: 1KZ7) show a contact between the DH-PH domain interface and the switch II region (red) of the respective substrate GTPase. **B**, a closer view of this contact shows a polar interaction between a switch II arginine and aspartate (red) and three conserved residues of the GEF (grey), two of which belong to the PH domain (Tyr889 and Gln834). **C**, Alignment of the predicted structure of the human obscurin DH-PH to the Dbs RhoA/Cdc42 complex shows that the corresponding obscurin residues are in a similar position as in the Dbs/substrate GTPase complexes. **D**, Guanosine nucleotide exchange factor activity of chimeric and zebrafish obscurin RhoGEF fragments towards RhoA, Cdc42, Rac1 and RhoQ/TC10. Black arrows indicate addition of buffer/GEF/EDTA. Data represent mean of n = 2-3 experiments. See Figure 2 for controls (experiments have been performed simultaneously in a multi-well plate reader).

This suggests the catalytic contribution of PH-domain residues is evolutionary conserved both in the trio-subfamily and across species. We thus hypothesized that these residues may need to be present for obscurin to exert GEF activity towards Rho GTPases. Since we were unable to purify any proteins containing the human obscurin PH domain, we cloned and attempted to purify several chimeric obscurin DH-PH fragments containing the human DH domain followed by the PH domain from human dbs and trio and the obscurin PH domain from chicken or zebrafish, all of which feature the aforementioned residues involved in supporting GEF activity. Of these chimeric proteins, only human/zebrafish chimeras (henceforth called “mermaid” obscurin) could be successfully purified (**Figure S8**). Many zebrafish RhoGTPases (in particular the putative substrate RhoA) have >90% sequence identity with human RhoGTPases ^23^. Given the high sequence identity of the substrates across species, we reasoned that zebrafish obscurin GEF fragments should likely exhibit activity towards human RhoGTPases.

We therefore cloned and successfully purified additional mermaid as well as several zebrafish obscurin DH and DH-PH fragments (**Figure S9**). Next, we tested mermaid and zebrafish RhoGEF fragments for nucleotide exchange activity towards RhoA, Cdc42, Rac1 and RhoQ/TC10. Again, we found no increase in the nucleotide exchange rate after addition of these GEF domains (**Figure 3D**). We also tested the nucleotide exchange activity of zebrafish obscurin DH-PH_5884−6217_ towards RhoB, RhoC, Rac2, Rac3 and RhoG but did not detect any increase in the nucleotide exchange rate (**Figure S10**).

Our data show, therefore, that the inclusion of the PH domain was not sufficient to obtain catalytically active obscurin RhoGEF *in vitro*, at least for the tested substrates.

### Obscurin RhoGEF domains can be phosphorylated by MST kinases and CaMKs

The complete absence of guanosine nucleotide exchange activity of DH-PH fragments is peculiar. Flanking SH3 and PH domains as well as linker regions are known to sometimes inhibit the activity of the DH domain ^24–28^. Typically, release of such autoinhibitory regions is mediated by phosphorylation (reviewed e.g. in Hodge and Ridley, 2016; Rossman et al., 2005). In the case of obscurin, simple autoinhibition of the obscurin DH domain by the SH3 or PH domain as the sole explanation for the lack of obscurin GEF activity is unlikely given the absence of activity of the isolated DH domain alone and the conserved positive catalytic contribution of the PH domain in trio-subfamily RhoGEFs. Interestingly, however, the murine obscurin RhoGEF region, too, has been reported to become phosphorylated at the residues corresponding to human Ser5669 (in the SH3-DH interdomain linker) and Thr5798 (within the DH domain) upon muscle exercise ^31^. Ser5669 is conserved among human, mice and zebrafish, Thr5798 only among human and mice (**Figure S11**). We thus speculated that phosphorylation might still be important to activate obscurin RhoGEF function, possibly via conformational changes in one of its domains.

To identify kinases that can phosphorylate the RhoGEF region, we used the largest human RhoGEF fragment available to us (SH3-DH) as a substrate in a commercial screen comprising a library of 245 Ser/Thr-protein kinases. Of these, 42 kinases exhibited a significant activity towards obscurin SH3-DH. Interestingly, the list of significant hits contained many CaMK-family kinases (CAMK1D, CAMK2B, CAMK2D and CAMK4) and MST-family kinases (MST1, MST2 and MST4), which also exhibited high total phosphorylation levels of obscurin SH3-DH (**Figure S12A**). A follow-up assay using the three kinases with the highest activity ratios (TBK1, CaMK4 and MST2) confirmed the activity (**Figure S12B**). While the relevance of CaMKs to cardiovascular biology is well documented, MST2 has been implicated e.g. in cardiac hypertrophic signalling in response to pressure overload ^32^.

To independently validate the phosphorylation of obscurin and narrow down the phosphorylation sites, we analysed phosphorylation of different GEF fragments (SH3, SH3-DH and DH) after addition of PKA, CaMK1d, CaMKII or MST2 using a phospho-protein specific staining method. In contrast to the screen, we found that only MST2 can robustly phosphorylate obscurin SH3-DH (**Figure 4A**). Interestingly, CaMKII did not phosphorylate the SH3-DH fragment but only the isolated SH3 domain, suggesting that the phosphorylation site is sterically inaccessible in the SH3-DH fragment, potentially due to an intramolecular interaction between SH3 and DH domain. None of the four tested kinases could phosphorylate the DH domain.

**Figure 4.**
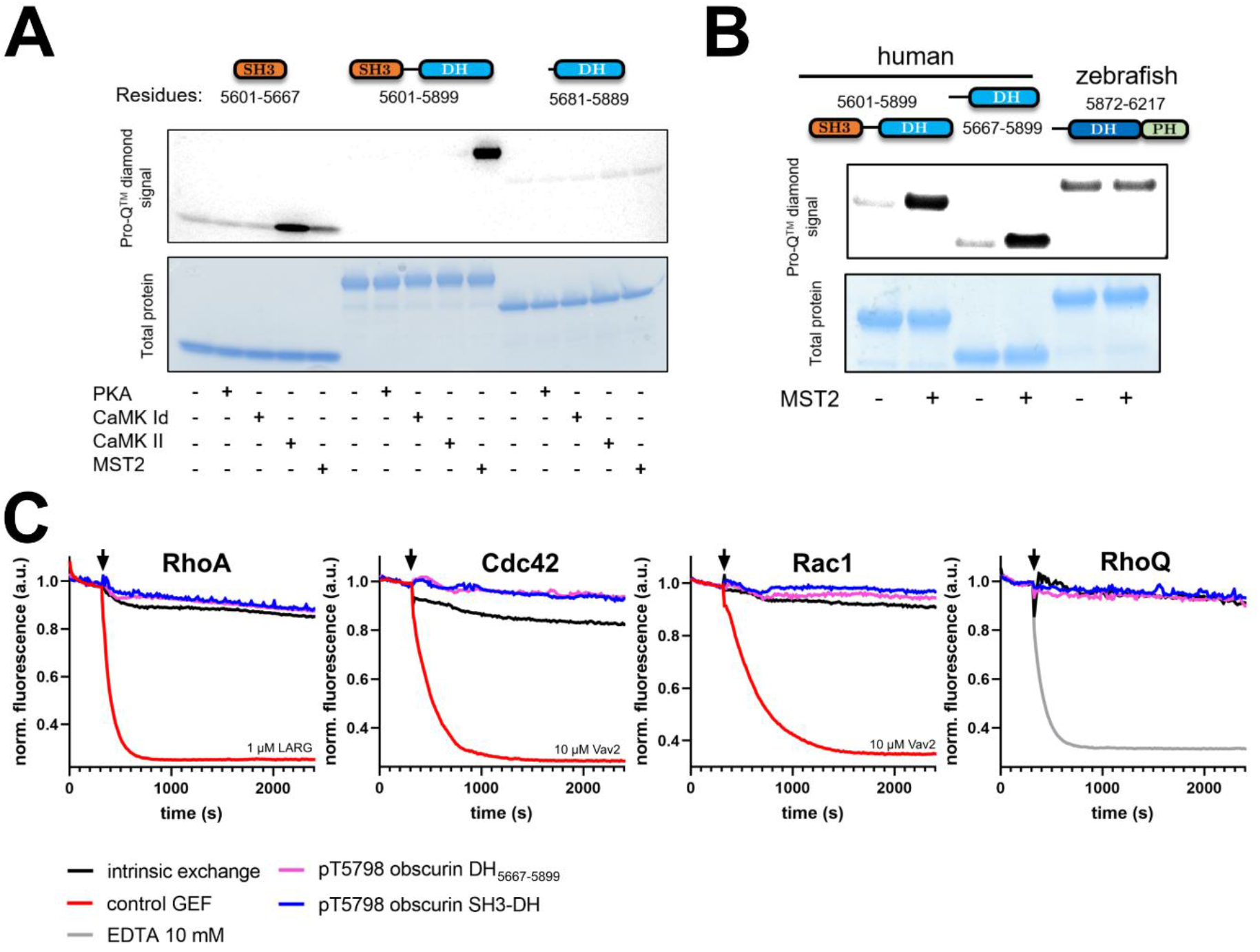
**A**, phosphorylation of different human obscurin RhoGEF domain combinations by different kinases. **B**, MST2-phosphorylation of an N- and C-terminally extended human DH domain and a zebrafish DH-PH fragment compared to human SH3-DH. **C**, Guanosine nucleotide exchange factor activity of obscurin RhoGEF (10 μM) fragments phosphorylated at Thr5798 towards RhoA, Cdc42, Rac1 and RhoQ/TC10. Black arrows indicate addition of buffer/GEF/EDTA. Data represent mean of n = 2-4 experiments.

We further tested whether phosphatases PP1 and PP2A, both of which are crucial regulators of myofilament and SR proteins, can dephosphorylate obscurin SH3-DH following phosphorylation by MST2. We found that both PP1 and PP2A can dephosphorylate the SH3-DH fragment (**Figure S12C**).

Next, we tested whether MST2 can phosphorylate the zebrafish DH-PH fragment and a human N- and C-terminally slightly extended DH domain featuring most of the interdomain linkers including the Ser5669 site. We found that MST2 can phosphorylate the extended human DH domain, but not the zebrafish DH-PH fragment (**Figure 4B**). Finally, we directly mapped the MST2 phosphorylation site using mass-spectrometry and confirmed that it is likely Thr5798 (**Figure S13**).

Having identified a kinase that phosphorylates the DH domain at a phosphorylation site reported in physiological context, we tested whether this modification leads to the activation of obscurin GEF activity by repeating the GEF activity assays with obscurin that had been phosphorylated *in vitro* by MST2. We found that phosphorylation of either SH3-DH or the N-terminally extended DH domain did not lead to discernible GEF activity towards RhoA, Cdc42, Rac1 or RhoQ/TC10 (**Figure 4C**).

Given the lack of activity *in vitro*, we wondered whether the obscurin DH-PH tandem could change the localization of endogenous RhoA and RhoQ, which might indicate GEF activity in cells. We thus overexpressed the GFP-tagged human obscurin DH-PH domains in neonatal rat and adult murine cardiomyocytes using adenoviral vectors and studied the localization of RhoA and RhoQ using immunofluorescence microscopy (**Figure S14**). In uninfected neonatal rat cardiomyocytes, RhoQ/TC10 exhibited a striated, RhoA a mostly diffuse localisation pattern (although very rarely also a striated pattern). RhoQ/TC10 and RhoA localisation in the obscurin DH-PH overexpressing cells did not appear any different to the cells expressing GFP alone, both in the presence and absence of the adrenergic agonist phenylephrine. Thus, we found no evidence for activity of obscurin RhoGEF domains in living cells. However, we cannot rule out the possibility that obscurin might activate RhoA or RhoQ/TC10 without changing their localisation, or that under different culture conditions or by using alternative cell types a difference would be revealed, warranting further work to test these possibilities.

## 3. Discussion

Although trio-subfamily GEFs are known for their reliance on the PH-domain for effective nucleotide exchange (Rossman and Campbell, 2000; Snyder et al., 2002), the complete absence of any catalytic activity of the recombinant obscurin GEF domains towards any of the 10 tested GTPases is surprising, given the previously reported activity of obscurin towards RhoA and RhoQ/TC10 using co-IP/pulldown assays ^12,13^. The putative activating effect of obscurin on RhoA seems to be conserved for the invertebrate homologues unc89 and Rho-1 ^33^. The inclusion of the PH domain or Thr5798 phosphorylation in vertebrate obscurin did not lead to activation of its RhoGEF function either. However, when used at sufficiently high concentrations such as in our study, even inefficient DH domains typically show some degree of discernible activity towards their substrate GTPases ^22^.

There are multiple reasons which could account for this discrepancy. Firstly, while our experiments tested many combinations of GEF fragments and RhoGTPases, they were, due to practical limitations, not exhaustive. Thus, we might have just missed the right GEF/substrate pair which would lead to discernible GEF activity (e.g. pThr5798-SH3-DH and RhoG). However, in a systematic kinetic study of 21 DH-domain RhoGEFs, all of the tested GEFs exhibited at least some discernible residual GEF activity towards RhoA, Rac1, or Cdc42, even when these were not the main physiological substrates ^18^. Since our minimal tested set of potential substrate GTPases included RhoA, Rac1, Cdc42 and RhoQ/TC10, we consider this possibility unlikely.

Another possibility is that the use of zebrafish obscurin or human/zebrafish chimeras might not work due to critical differences in the sequence and mechanism between species. Human and zebrafish RhoA exhibit a very high (>90%) sequence identity ^23^ and zebrafish obscurin PH residues known to be important in the GTPase/PH-domain interaction, too, are conserved as shown above. The conservation of molecular function is a feature often observed in the evolution of Ras-family GTPases. Yeast Ypt1, for instance, can be substituted by its mouse homologue without loss of function ^34^. For RhoGTPases in particular, evolutionary diversification of function rather occurs via diversification of their ‘regulatome’ ^21^. For these reasons, we think species differences are not the most likely explanation for the lack of observed activity either.

A more likely explanation, in our view, is that the activation of obscurin requires other, potentially even multiple factors to obtain an active GEF. For instance, we do not yet know what effect phosphorylation of Ser5669 has on obscurin function or what the responsible kinase is for this site. Perhaps activation of obscurin RhoGEF function even requires both DH and PH domains plus phosphorylation at either or both Ser5669 and Thr5798 or phosphorylation-dependent recruitment of additional cellular cofactors. We conducted molecular dynamics simulations using a structural model of the SH3-DH-PH domains that suggested that phosphorylation at Ser5669, Thr5798, or at both positions does not lead to large conformational changes, inter-domain motions or significantly altered dynamics for individual residues (**Figure S15**). Thus, phosphorylation at these sites might change the availability of bindings sites for interaction partners rather than altering intramolecular conformation. Furthermore, it is worth pointing out that we used cardiomyocytes for cell biological experiments, whereas Ford-Speelman and colleagues used *tibialis anterior* muscles ^13^, suggesting that such additional factors required for GEF activity might even be tissue specific. It is interesting to note in this context that the complete knockout of obscurin in mice leads to a fairly mild muscle phenotype with disrupted organisation of the sarcoplasmic reticulum and altered cellular calcium-handling^5,35^, while human loss-of-function mutations in OBSCN, abrogating the C-terminal GEF domain, display a compatible phenotype with susceptibility to severe rhabdomyolysis due to defective calcium-handling^36^. Neither phenotype offers an obvious mechanistic link to defective small GTPase signalling, other than potentially in membrane remodelling during SR-formation.

Another possibility to consider is that the obscurin RhoGEF domains might have lost its catalytic function during evolution. Enzymes which lost their catalytic function during evolution include proteins from various classes including GTPases, kinases and phosphatases ^37,38^. The N-terminal kinase of the *C. elegans* obscurin homologue unc89, for example, is predicted to be inactive due to the alteration of key residues involved in catalysis and the absence of the N-terminal kinase lobe ^39^. Such enzymes are often called pseudo-enzymes and obscurin might thus be the first instance of a “pseudo-GEF”. In this case, the observed effect of obscurin on RhoGTPase activity^12,13^ could be indirect. Obscurin might, for example via a non-catalytic interaction, recruit GTPases to specific membrane compartments where they could be subsequently activated by another GEF.

A more detailed bioinformatic analysis of the trio-subfamily DH-PH domain sequences suggests that obscurin does not cluster with other trio-subfamily members and shows the least similarity to other subfamily members (**Figure S16A**). Furthermore, obscurin exhibits several differences at multiple highly conserved and functionally important residues for the interaction with GTPases ^18^ (**Figure S16B**). These analyses, together with our experimental data, demonstrate that obscurin is a GEF that, if catalytically active, appears to be regulated in a more complex fashion than other RhoGEFs. Clearly, these results warrant further investigation into the activity of the obscurin DH-PH domains in living cells. A promising approach would be to use FRET-biosensors, which are available for both RhoA and RhoQ/TC10 (as well as many other RhoGTPases) to monitor the activity of RhoA or RhoQ/TC10 in response to obscurin in real-time ^40,41^.

Despite the lack of observed catalytic activity, the availability of highly pure recombinant protein fragments paves the way for the identification and structural or biophysical characterization of further interaction partners. Moreover, the identification of multiple kinases such as CaMKs and MST-kinases as potential regulators of obscurin RhoGEF function opens new avenues to study the role of these domains in cells.

## 4. Material and Methods

Details on material and methods, including a list primers used for cloning (**Table S1**), can be found in the online supplementary information.

## Author Contributions

DK, MR and MG designed the experiments. DK, ALK and AF performed experiments and analysed the data. AA developed the small-scale purification assay. AB and MR performed the molecular dynamics simulations. MP performed the NMR experiments. MP, MR and MG supervised the study. DK wrote the paper with help from MR and ALK. MG conceived the study and acquired funding. All authors proofread the manuscript.

## Acknowledgements

We thank Dr Thomas Kampourakis for access to the plate reader. DK was funded by a PhD studentship from the British Heart Foundation (grant [FS/17/65/33481]). This work was supported by funds from the Wellcome Trust (Collaborative Award in Sciences 201543/Z/16/Z to MG), the European Research Council under the European Union’s Horizon 2020 Programme (ERC-2019-SyG, grant no. 856118 to MR and MG) and the Medical Research Council (MR/R003106/1 to MG and ALK). Earlier work leading to the present paper was supported by the Volkswagen Foundation Research grant I/81 797 and I/78 989 to MG. MG holds the BHF Chair of Molecular Cardiology. We acknowledge support by CREATE^42^. The authors declare that they have no conflict of interest.

