## Supplemental Information for "Obscurin Rho GEF domains are phosphorylated by MST-family kinases but do not exhibit nucleotide exchange factor activity towards Rho GTPases *in vitro*"

#### Material and Methods

*Cloning and viral packaging*

Human and chimeric obscurin GEF constructs 1-45 (cf. Figures S2 and S8) have been generated with standard PCR techniques using the primers listed in Table S1 followed by transfer of PCR products into a modified pET15b vector (Novagen) with an N-terminal hexahistidine-tag and a TEV cleavage site using the NEBuilder® HiFi DNA Assembly kit (NEB, cat. no. E2621S). C-terminally truncated (for removal of prenylation site) small GTPases $RhoA_{1\text{-}181,F25N}$, $RhoB_{1\text{-}188,F25N}$, $RhoC_{1\text{-}181,F25N}$ ,$Rac1_{1\text{-}177}$, $Rac2_{1\text{-}177}$, $Rac3_{1\text{-}177}$, $RhoG_{1\text{-}177}$, $Cdc42_{1\text{-}178}$ and $RhoQ/TC10_{1\text{-}185}$ were cloned into a modified pGEX-2TK vector (GE Healthcare) in which the thrombin cleave-site was substituted by a TEV cleavage site. Cloning of constructs 46-58 (cf. Figure S9) and larg and vav2 DH domains was done by Bio Basic Inc (Canada, https://www.biobasic.com/). Cloning and viral packing of N-terminally eGFP-tagged human obscurin DH-PH (residues 5681-6019, numbering as in human obscurin B, NCBI ref. seq. NM_001098623) into either AAV9 or AdV vectors was performed by VectorBuilder Inc (United States, https://en.vectorbuilder.com/). All constructs were validated by sequencing.

*Protein expression and purification*

For protein expression, BL21-CodonPlus (DE3)-RIPL (Agilent, cat. no. 230280) *E.coli* cells were heat-shock transformed with expression vectors and grown over night on LB agar plates with carbenicillin. The next morning, 0.5 to 2L LB + carbenicillin were inoculated with cells scraped directly from the plate and grown at 37°C in a shaking incubator at 180-200 rpm until the culture reached an optical density at 600 nm (OD600nm) of 0.6 - 0.9, at which point expression was induced by addition of 0.2 mM IPTG and allowed to proceed overnight at 20°C. Cells were harvested the following morning and lysed via 2x freeze/thaw cycles followed by addition of lysozyme into the following lysis buffer: 50 mM Tris-HCl pH 7.5, 300 mM NaCl, 25 mM imidazole, 1:1000 (v/v) β-mercaptoethanol, (5mM $MgCl_2$ for GTPases), cOmplete™ EDTA-free Protease Inhibitor Cocktail (Roche, cat. no. 11873580001).
RhoGTPases were purified using a GSTrap™ 4B column (GE Healthcare, cat. no. 29048609) followed by size exclusion chromatography into a final buffer of 50 mM Tris-HCl, 100 mM NaCl, 2 mM $MgCl_2$ and 2 mM DTT. Obscurin SH3, SH3-DH, $DH_{5681\text{-}5889}$ and $DH_{5667\text{-}5899}$ fragments were purified using a HisTrap™ FF column (GE Healthcare, cat. no. 29048609) followed by size exclusion chromatography into a final buffer of 30 mM Hepes pH 7.5, 100 mM NaCl and 2 mM DTT. Chimeric and zebrafish obscurin fragments, vav2 and larg DH domains were batch-purified using a single step affinity enrichment with Ni-NTA resin (Qiagen, cat. no. 30410) followed by 3x washing in lysis buffer before proteins were eluted from the resin and buffer exchanged into final buffer (30 mM Hepes pH 7.5, 100 mM NaCl and 2 mM DTT). Following purification, each protein was aliquoted, flash-frozen in liquid $N_2$ and stored at -70°C until further use.
For small scale test expression and purification for the purpose of solubility screening, cells were grown for 3-4 days in 4 mL autoinduction medium (Studier, 2005) supplemented with carbenicillin in 24 deep-well plates (Elkay, cat. no. 43001-0066). Following transformation, cells were grown for ~6hrs at 37°C before the temperature was lowered to 20°C and cells were grown until harvesting. Expression of proteins was monitored using cells transformed with GFP or mCherry as indicator cultures. After pelleting, cells were lysed in the plates using Bacterial Protein Extraction Reagent (Thermo Scientific, cat. no. 78243) and purified with 200 µl of Ni-NTA resin (Qiagen, cat. no. 30410). After two washing steps (buffer A: 50mM Tris-HCl pH 7.7, 300 mM NaCl, 2 mM DTT, 30 mM imidazole), the protein was eluted with a volume of 2x400 µL elution buffer (like buffer A but with 1M imidazole).
Protein concentration was determined either by absorbance at a wavelength of 280 nm using a DS-11 FX+ Spectrophoto-/Fluorometer (DeNovix) or for GTPases, due to interference of the guanosine nucleotide, by the Bradford assay (VWR Life Sciences, cat. no. M172-1L) using BSA as a standard.

**Table S1**: Primers used in this study

| Primer | Sequence | Tm (°C) | Comments |
|---|---|---|---|
| 4 | CACGCGTGGATCCTTActggcggatgggct c | 66 | Obscurin 5899 reverse, 5' overlap with pET6HtevC2 |
| 7 | TATTTTCAGGGCTCGAGCaagctgtcacc tgagt gg | 64 | Obscurin 5667 forward, 5' overlap with pET6HtevC2 |
| 8 | TATTTTCAGGGCTCGAGCcctggggagg ctg | 62 | Obscurin 5681 forward, 5' overlap with pET6HtevC2 |
| 9 | TATTTTCAGGGCTCGAGCtctgaagacga atac aaggc | 62 | Obscurin 5686 forward, 5' overlap with pET6HtevC2 |
| 10 | TATTTTCAGGGCTCGAGCgaatacaagg caag gctgag | 63 | Obscurin 5689 forward, 5' overlap with pET6HtevC2 |
| 11 | TATTTTCAGGGCTCGAGCaaggcaaggc tgagctc | 65 | Obscurin 5691 forward, 5' overlap with pET6HtevC2 |
| 12 | CACGCGTGGATCCTTActccatgagggac acgtg | 65 | Obscurin 5884 reverse, 5' overlap with pET6HtevC2 |
| 13 | CACGCGTGGATCCTTAggtgcctgggtagtt ctc | 65 | Obscurin 5889 reverse, 5' overlap with pET6HtevC2 |
| 14 | CACGCGTGGATCCTTActgcagggtgcctg | 63 | Obscurin 5891 reverse, 5' overlap with pET6HtevC2 |
| 15 | CACGCGTGGATCCTTActcgcccagggc | 62 | Obscurin 5895 reverse, 5' overlap with pET6HtevC2 |
| 16 | TATTTTCAGGGCTCGAGCgagcccatccg | 63 | Obscurin 5895 forward, 5' overlap with pET6HtevC2 |
| 17 | TATTTTCAGGGCTCGAGCcacttcatcgtg tgg | 63 | Obscurin 5901 forward, 5' overlap with pET6HtevC2 |
| 18 | CACGCGTGGATCCTTAacgctgctggatgc | 62 | Obscurin 6005 reverse, 5' overlap with pET6HtevC2 |
| 19 | CACGCGTGGATCCTTAaggcagggccag ac | 64 | Obscurin 6009 reverse, 5' overlap with pET6HtevC2 |
| 20 | CACGCGTGGATCCTTAccgccacacaggc | 64 | Obscurin 6012 reverse, 5' overlap with pET6HtevC2 |
| 21 | tgggtagttctccatgagg | 63 | Obscurin 5887 reverse |
| 22 | ctccatgagggacacg | 62 | Obscurin 5884 reverse |
| 23 | ctcgcccagggc | 62 | Obscurin 5895 reverse |
| 24 | CGTGTCCCTCATGGAGggctatgacggga atctc | 62 | Dbs 787 forward, 5' overlap with human obscurin |
| 25 | CACGCGTGGATCCTTAttctctacaagcctg cagc | 64 | Dbs 921 reverse, 5' overlap with pET6HtevC2 |
| 26 | CGTGTCCCTCATGGAGgggtttgatgaaaa cattgag | 63 | Trio 1475 forward, 5' overlap with human obscurin |
| 27 | CAGGCCCTGGGCGAGctcatcctacagga atc | 64 | Trio 1486 forward, 5' overlap with human obscurin |
| 28 | CACGCGTGGATCCTTAcgtccgctcctgga tg | 65 | Trio 1594 reverse, 5' overlap with pET6HtevC2 |
| 29 | CTCATGGAGAACTACCCAgg | 64 | Obscurin (chicken) 7963 forward, 5' overlap with human obscurin |
| 30 | CACGCGTGGATCCTTAgtctggagggatcc agac | 64 | Obscurin (chicken) 8096 reverse, 5' overlap with pET6HtevC2 |
| 31 | CTCATGGAGAACTACCCAgccaatc | 69 | Obscurin (zebrafish) 6090 forward, 5' overlap with human obscurin |
| 32 | CACGCGTGGATCCTTAatctggagagcac catgttg | 64 | Obscurin (zebrafish) 6217 reverse, 5' overlap with pET6HtevC2 |

*Dot-blotting*

For analysis of protein solubility, 1-2 µl samples from the purification screen were applied to a nitrocellulose membrane and allowed to dry for 1-2 hrs before the membrane was blocked for 15-30 min in with 5% (w/v) milk powder in antibody binding buffer (10 mM Tris pH 7.4, 9 g/l NaCl, 1 ‰ (v/v) Tween-20). Mouse anti-His mAb (Millipore, cat. no. 70796) was applied at a dilution of 1:1000 for 1hr at RT or overnight at 4°C. Secondary antibodies (HRP-conjugated rabbit anti-mouse IgG, Dako, cat. no. P0260) were applied at a dilution of 1:1000 for 1 h at room temperature before signals were detected by chemiluminescence on a ChemiDocTM XRS+ imaging system (Bio-Rad) using Clarity ECL Western Substrate (Bio-Rad, cat. no. 1705061).

*Buffer exchange*

To exchange buffer of proteins, the following desalting columns appropriate for the sample volume were used according to manufacturer's instructions: 30-130 µL sample volume with Zeba™ Spin Desalting Columns, 7K MWCO (Thermo Scientific™, cat. no. 89882) or illustra™ NAP-5/10/25 columns for sample volumes of 0.5/1/2.5 mL (GE Healthcare, cat. nos. 17-0853-01/17-0854-01/17-0852-01).

*1D-NMR*

We used 1D-NMR to analyse selected protein fragments and assess whether the resulting NMR spectra exhibit features that indicate correct folding of the domain (Kwan et al., 2011). The protein to be analysed was buffer exchanged into 25 mM $HNa_2PO_4$ pH 7.3, 100 mM NaCl, 2 mM DTT. The protein was concentrated to at least 60 µM and spectra were recorded with an Ascend 600 instrument (Bruker Scientific Instruments).

*Nucleotide exchange kinetics*

Nucleotide exchange kinetics of RhoGTPases were measured as described previously (Itzen et al., 2007; Koch et al., 2016). Briefly, GTPases were preparatively pre-loaded with fluorescent mant-GDP by adjusting the protein concentration to 50-100 µM and spiking the sample with a 5-fold molar excess of mant-GDP (Jena Bioscience, cat. no. NU-204S) and 5 mM EDTA. After incubation of 2-4 hrs at RT in a light-protected tube, the buffer was exchanged to 50 mM Tris-HCl, 100 mM NaCl, 2 mM $MgCl_2$ and 2 mM DTT and the protein was aliquoted, flash-frozen in liquid $N_2$ and stored at -70°C until further use.

Nucleotide exchange reactions were recorded on a CLARIOstar microplate reader (BMG LABTECH, Germany) in 96-well microplates (greiner bio-one, cat. no. 675076 or invitrogen, cat. no. M33089) at 25°C. At the beginning of each experiment, 80 µL of 1.875 µM [GTPase] in reaction buffer equilibrated to room temperature (40 mM Hepes pH 7.5, 100 mM NaCl, 5 mM $MgCl_2$, 2 mM DTT) were pipetted into the plate and the signal was recorded (excitation wavelength $\lambda_{ex}$ = 360 nm, measured emission wavelength $\lambda_{ex}$ = 440 nm, sampling every 2-15s) for a sufficiently long period to reach a stable baseline (about 300 s). The nucleotide exchange reaction was initiated by addition of 20 µL of GDP or GDP + EDTA or GDP + GEF in buffer so that the final concentrations were 1.5 µM for the GTPase, 200 µM GDP, 10mM EDTA, or 0.5 - 25 µM GEF. After mixing by careful pipetting, the signal was recorded for further 1 - 2 hrs. The obtained traces were normalised by the average fluorescence signal of the first 300 s before start of the reaction and apparent rate constants $k_{obs}$ were obtained from the normalised traces by fitting the decay phase of the signal to a mono-exponential decay function of the form $f(t) = (a_0 - a_{plateau})\ e^{-kobs\ x\ t} + a_{plateau}$ in the GraphPad Prism software v8.3 (GraphPad Software, Inc).

*Kinase screening*

Kinase screening using the obscurin SH3-DH domain as a substrate was performed externally with the KinaseFinder screen from ProQinase GmbH (now Reaction Biology Europe GmbH, https://www.reactionbiology.com/). Details on the procedure provided by ProQinase can be found in Appendix A.

*Phosphorylation of obscurin*

For analytical purposes, 0.2 mg/ml obscurin RhoGEF fragments (5-20 µM, depending on the fragment used) were phosphorylated by 200 U PKA (Sigma-Aldrich, cat. no. 539576 ) or 500 U CaMK II (NEB, cat. no. P6060L) or 25 ng CaMK Id (BioVision, cat. no. 7713-5) or 25 ng MST2 (Millipore, cat. no. 14-524) in kinase reaction buffer (30 mM Hepes pH 7.5, 100 mM NaCl, 2 mM $MgCl_2$, 300 µM ATP, 2 mM DTT [+1mM $CaCl_2$ and 1 µg/ml (approx. 3 µM) Calmodulin (NEB, cat. no. 6060S) for CaMK II/Id]). Samples were incubated for 3 h at 30°C in a PCR cycler and the reaction was stopped by addition of SDS sample buffer. For visualisation of phosphoprotein, 2 µg of obscurin substrate were separated by SDS-PAGE and stained with Pro-Q™ Diamond Phosphoprotein Gel Stain as described below. For preparative MST2-phosphorylation of obscurin SH3-DH or $DH_{5667-5899}$, 500 ng MST2 were added to a reaction volume of 200 µL kinase reaction buffer containing a substrate concentration of 110 µM and the reaction was allowed to proceed for 5 hrs at 25°C, followed by further incubation overnight at 8°C. The phosphorylated protein was buffer exchanged into 40 mM Hepes pH 7.5, 100 mM NaCl and 2 mM DTT.

*Dephosphorylation of MST2-phosphorylated obscurin*

Following preparative phosphorylation of obscurin SH3-DH by MST2, 0.2 mg/ml obscurin phosphosubstrate were incubated with 1 µM of phosphatases PP1 or PP2A (Cayman Chemical, cat. no. 10011237) or 1 U rAPid Alkaline Phosphatase (Roche, cat. no. 4898133001) in 40 mM Hepes pH 7.5, 100 mM NaCl, 5mM $MgCl_2$ and 2 mM DTT (+ 1mM $MnCl_2$ for activation of PP1) in a reaction volume of 50 µL. The reaction was allowed to proceed for 1.5 h at 25°C and was stopped by the addition of SDS sample buffer.

*Phosphoprotein staining*

For detection of protein phosphorylation on polyacrylamide gels, Pro-Q™ Diamond Phosphoprotein Gel Stain (Invitrogen, cat. no. P33301) was used according to the manufacturer's instructions.

*Mass-spectometry*

The MST2 phosphorylation site in the SH3-DH fragment was identified externally using a commercial mass-spectometry service from the Metabolomics and Proteomics Laboratory of the Bioscience Technology Facility of the University of York. Details on the procedure can be found in Appendix B.

*Neonatal ventricular rat cardiomyocyte preparation*

Neonatal ventricular rat cardiomyocytes (NRCs) were isolated from Wistar rat pups and cultured as described previously (Simpson and Savion, 1982). In brief, hearts were isolated from Wistar rat pups at postnatal day 0 to 2 and cut into 4 in ice cold ADS (116 mM NaCl, 20 mM HEPES, 0.8 mM $NaH2PO_4$, 5.6 mM glucose, 5.4 mM KCl, 0.8mM $MgSO_4$ pH 7.35). The hearts were enzymatically digested in a sequential manner by incubation in enzyme solution containing collagenase type II (Worthington) (57.5 U/ml) and pancreatin (Sigma) (1.5 mg/ml) for 4-5 times for 15 min in a shaking incubator at 37°C. The supernatant is collected into medium containing 5% FCS and passed through a 70 micron cell strainer (Falcon Corning) before being pelleted at low speed. The cells were pre-plated onto 90 mm dishes (Nunc) in plating medium (DMEM, 5% FCS, 10% HS, non-essential amino acids, penicillin/streptomycin (P/S) and L-glutamine) for 2 h to allow non-myocytes to adhere. The non-adherent cardiomyocyte enriched fraction is then plated onto collagen (Attachin, Genlantis) coated 35mm dishes (Nunc) and cultured at 37°C and 5% $CO_2$. Once the cells had recovered (2-3 days), the non-adherent cells were washed away with culture medium (M199, DBSSK [116mM NaCl, 1 mM $NaH2PO_4$, 0.8 mM $MgSO_4$, 32.1 mM $NaHCO_3$, 5.5 mM glucose, 1.8 mM $CaCl_2$ pH7.2], 4% Horse serum, P/S and L-glutamine) and cultured until day 8-9 for further maturation (medium exchange every 3 days). The NRCs were transduced with 1µl of AdGFP or AdDHPH (at $1.05 \times 10^{11}$ and $9.16 \times 10^{10}$ IFU/ml respectively) for 24 hours then cultured in the absence and presence of phenylephrine (100µM, Sigma) for another 24hours before being fixed with 4% paraformaldehyde for 10 mins.

*Immunostaining*

The cells were permeabilised with 50µg/ml digitonin and blocked with 10% goat serum in IF buffer (10% bovine serum albumin, 0.1mM Tris pH7.5, 15.5mM NaCl, 0.2mM EGTA, 0.2mM $MgCl_2$) for 30mins at room temperature before incubation with primary antibody (rabbit anti-rhoQ/TC10 Abcam, cat. no. ab32079, mouse anti-rhoA Sigma, cat. no. SAB140017, both 1:100, rabbit anti titin Z1Z2 1:100 (Gautel et al., 1996), mouse anti-myomesin b4 1:50 (Grove et al., 1984)) overnight at 4°C. The cells were then washed 3 times 5 mins in PBS, and incubated with secondary antibody (Cy3 anti mouse, JIR, cat. no. 115-165-146, Cy5 anti rabbit, JIR, cat. no. 111175144 111561, DAPI, Sigma, all 1:100) for 1 hr at room temperature. After washing 3x 5 mins with PBS, the dishes were mounted with mounting medium (30mM Tris pH 9.5, 0.24M n-propyl gallate, 70% glycerol) and a coverslip applied and sealed with clear nail varnish before being imaged on a Zeiss LSM 510 confocal microscope.

*Western Blotting*

HEK293 cells were transduced with 1µl of AdGFP or AdDHPH (at $1.05 \times 10^{11}$ and $9.16 \times 10^{10}$ IFU/ml respectively) for 24 hours. The cells were scraped off the dish and resuspended in SDS-loading buffer (Laemmli). Homogenate samples or recombinant GTPases were loaded onto a 4-15% acrylamide gel (Biorad). The proteins were transferred onto nitrocellulose overnight at 60mA and blocked with 10% nonfat dry milk in lo-salt buffer (0.9% w/v NaCl, 10mM Tris pH 7.4, 0.1% tween-20) and probed with anti-HIS (Millipore, cat. no. 70796), anti-GFP (Roche) or anti-obscurin DH antibody (Young et al., 2001) for 1 hour at RT. The blot was washed in lo-salt buffer 3 x 5 mins and incubated with HRP-tagged secondary antibodies (DAKO) for 1 hour. The blot was then washed again and signals were detected by chemiluminescence on a ChemiDocTM XRS+ imaging system (Bio-Rad) using Clarity ECL Western Substrate (Bio-Rad, cat. no. 1705061).

*Bioinformatic analyses*

Multiple sequence alignment of amino acid sequences was performed with clustal Omega server v1.2.4. (EMBLEBI, https://www.ebi.ac.uk/Tools/msa/clustalo/) (Sievers et al., 2011).

Aligned sequences and amino acid properties such as hydrophobicity or percent identity across aligned sequences) were visualised in the UGENE software v1.28.1 (Unipro) (Okonechnikov et al. 2012).

Sequence conservation across species species was assessed with ConSurf (https://consurf.tau.ac.il/) (Ashkenazy et al., 2010) with the following parameters: amino acids, no known structures, no MSA upload (sequence was provided in FASTA format), Proteins database: Uniprot, Select homologs for ConSurf analyses: automatically. Other parameters were left at their default settings.

Secondary structure predictions for proteins were obtained from 2018 to 2019 using the PredictProtein (https://predictprotein.org/) and PSIPRED 4.0 (http://bioinf.cs.ucl.ac.uk/psipred/) with default parameters (Buchan and Jones, 2019; Yachdav et al., 2014).

Homology based tertiary structure models were obtained with I-TASSER (https://zhanggroup.org/I-TASSER/) at default parameters (Roy et al., 2010) and visualised with PyMOL Molecular Graphics System v2.3.4 (https://pymol.org/2/, Schrödinger, LLC).

*Molecular dynamics simulations*

Before the simulations, a predicted structure of the obscurin SH3-DH-PH domain triplet was generated using the ColabFold AlphaFold2 server (Jumper et al., 2021; Mirdita et al., 2022) using default options with Amber relaxation and a protein sequence corresponding to residues 5602 to 6008 of obscurin transcript variant 1. Prior to molecular dynamic simulations, phosphorylated models were generated in Coot (Emsley et al., 2010) by either replacement of Ser5669 with phosphoserine, replacement of Thr5798 with phosphothreonine or replacement of both.

All-atom, solvated molecular dynamics simulations were run following energy minimisation for the four SH3-DH-PH models in Amber (Salomon-Ferrer et al., 2013) using the ff14SB forcefield with a sampling rate of 100ps, temperature of 298k, ionic strength of 0.1M and a simulation time of 1ms. The simulations were repeated 5 times for each model. Data including RMSD and RMSF was extracted from the molecular dynamic trajectories using scripts from ccptraj (Roe and Cheatham, 2013) and Bio3D (Grant et al., 2021).

### Supplementary figures

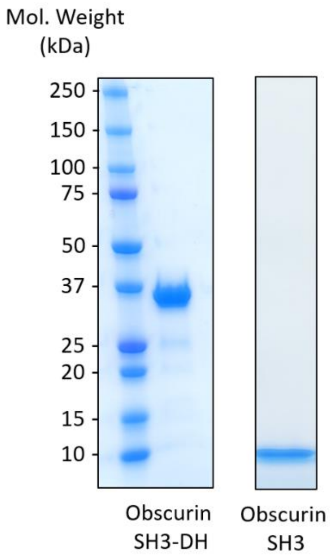


**Figure S1**. **A**, SDS-PAGE analysis of obscurin SH3-DH (amino acids 5601-5899) and SH3 domains (amino acids 5601-5667) after affinity purification and size-exclusion chromatography.

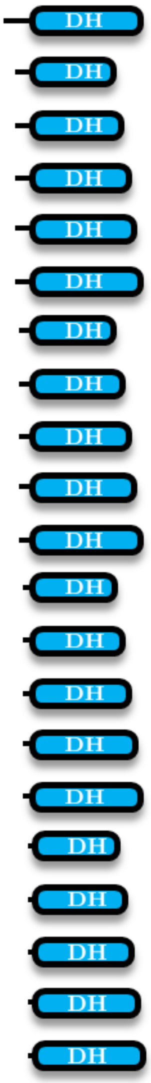


| Nr. | Primers | Residues (obscurin B) |
|---|---|---|
| 1 | 7,4 | 5667-5899 |
| 2 | 8,12 | 5681-5884 |
| 3 | 8,13 | 5681-5889 |
| 4 | 8,14 | 5681-5891 |
| 5 | 8,15 | 5681-5895 |
| 6 | 8,4 | 5681-5899 |
| 7 | 9,12 | 5686-5884 |
| 8 | 9,13 | 5686-5889 |
| 9 | 9,14 | 5686-5891 |
| 10 | 9,15 | 5686-5895 |
| 11 | 9,4 | 5686-5899 |
| 12 | 10,12 | 5689-5884 |
| 13 | 10,13 | 5689-5889 |
| 14 | 10,14 | 5689-5891 |
| 15 | 10,15 | 5689-5895 |
| 16 | 10,4 | 5689-5899 |
| 17 | 11,12 | 5691-5884 |
| 18 | 11,13 | 5691-5889 |
| 19 | 11,14 | 5691-5891 |
| 20 | 11,15 | 5691-5895 |
| 21 | 11,4 | 5691-5899 |

Dot-blot signal

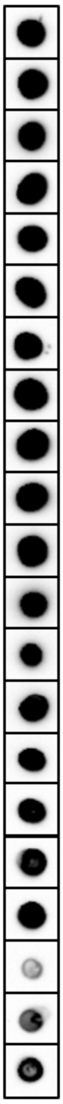

**Figure S2**. Construct design, primers used and dot-blot analysis of the eluted fraction after affinity purification of the human obscurin DH, DH-PH and PH domains.

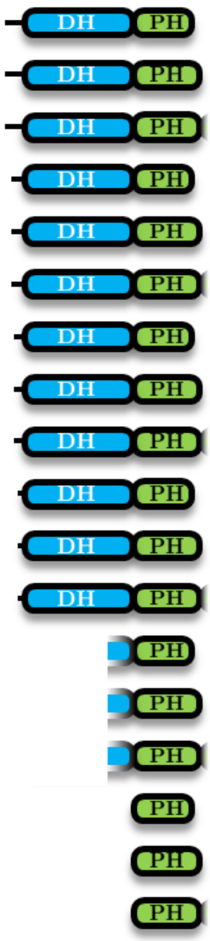

| Nr. | Primers | Residues (obscurin B) |
|---|---|---|
| 22 | 8,18 | 5681-6005 |
| 23 | 8,19 | 5681-6009 |
| 24 | 8,20 | 5681-6012 |
| 25 | 9,18 | 5686-6005 |
| 26 | 9,19 | 5686-6009 |
| 27 | 9,20 | 5686-6012 |
| 28 | 10,18 | 5689-6005 |
| 29 | 10,19 | 5689-6009 |
| 30 | 10,20 | 5689-6012 |
| 31 | 11,18 | 5691-6005 |
| 32 | 11,19 | 5691-6009 |
| 33 | 11,20 | 5691-6012 |
| 34 | 16,18 | 5895-6005 |
| 35 | 16,19 | 5895-6009 |
| 36 | 16,20 | 5895-6012 |
| 37 | 17,18 | 5901-6005 |
| 38 | 17,19 | 5901-6009 |
| 39 | 17,20 | 5901-6012 |

Dot-blot signal

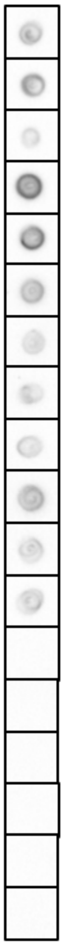

**Figure S2 (continued)**. Construct design, primers used and dot-blot analysis of the eluted fraction after affinity purification of the human obscurin DH, DH-PH and PH domains.

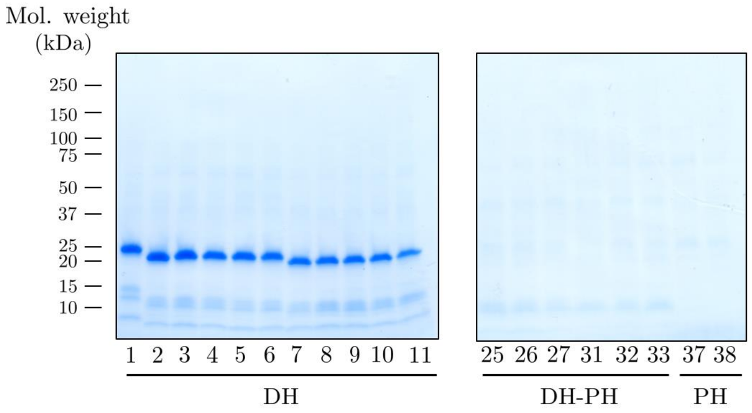


**Figure S3** SDS-PAGE analysis of selected fragments after affinity purification. The same samples shown in the dot-blots of Figure S2 were used for SDS-PAGE.

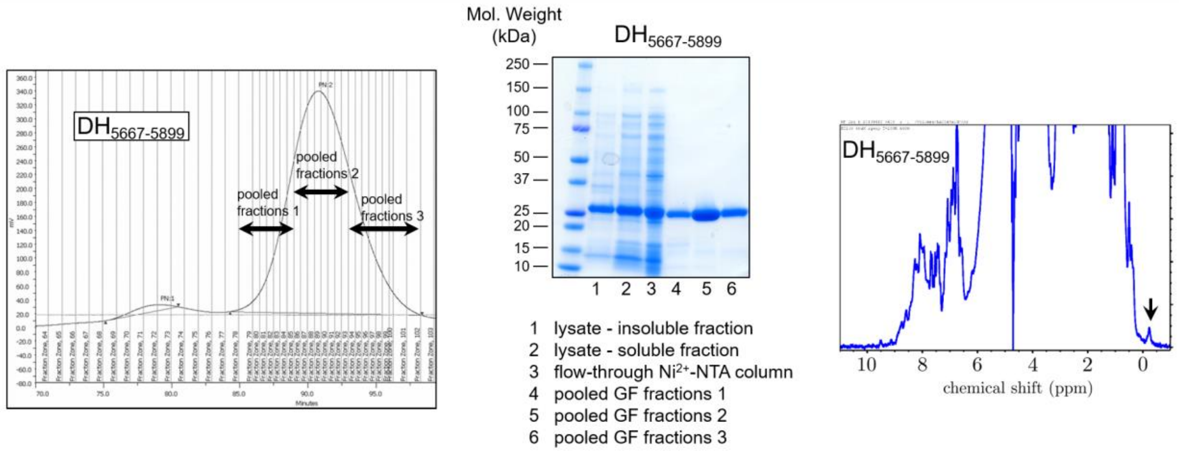


**Figure S4**. **Left**, gel-filtration elution profile of DH domain fragment comprising residues 5667-5899. **Middle**, SDS-PAGE analysis of samples at different steps of purification process of $DH_{5667\text{-}5899}$ fragment. **Right**, 1D-NMR analysis of purified $DH_{5667\text{-}5899}$ fragment. NMR spectrum exhibits wide peak dispersal and peaks below 0 ppm (black arrow), indicating that the protein is folded.

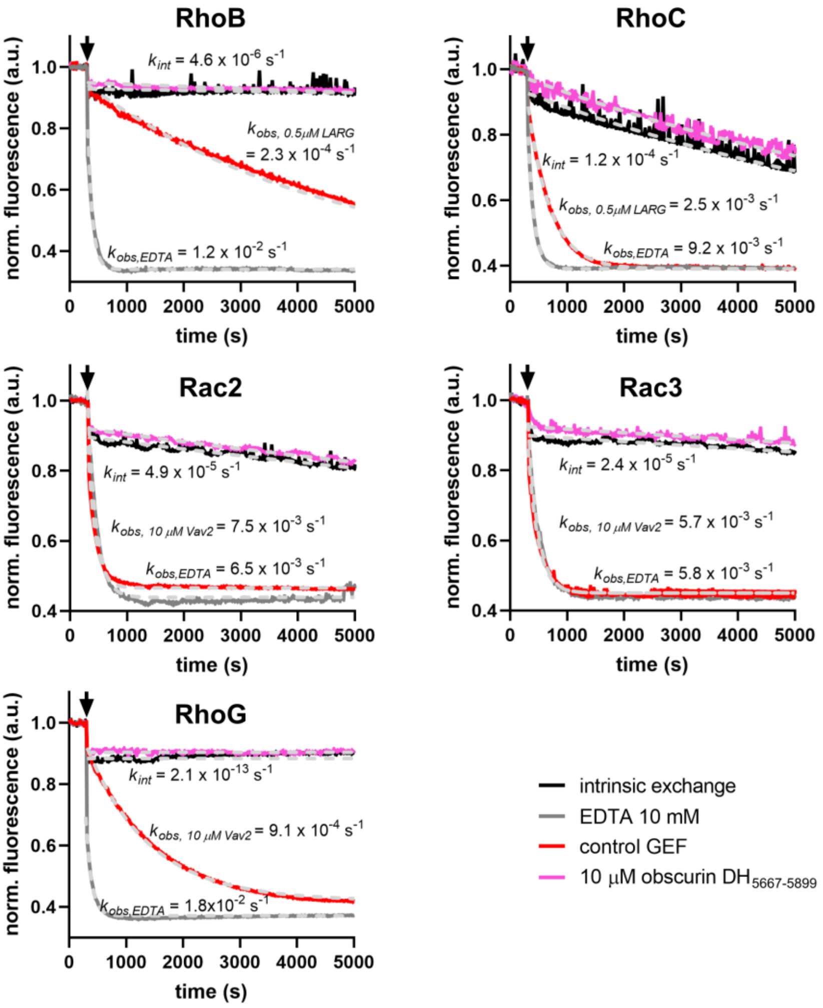


**Figure S5**. Guanosine nucleotide exchange factor activity of obscurin RhoGEF fragments towards RhoB, RhoC, Rac2, Rac3 and RhoG. Black arrows indicate addition of buffer/GEF/EDTA. Data represent mean of n = 2-3 experiments.

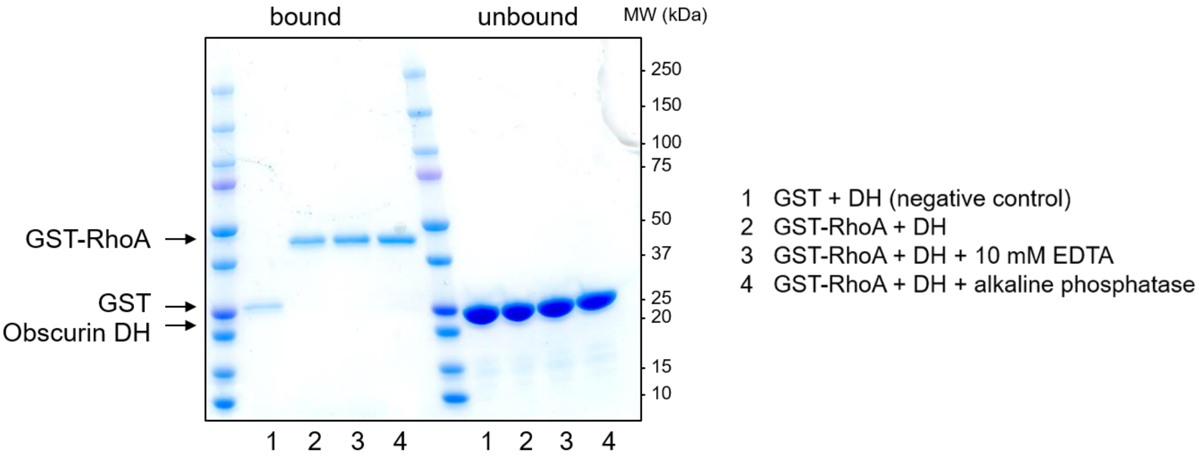


1 GST + DH (negative control)
2 GST-RhoA + DH
3 GST-RhoA + DH + 10 mM EDTA
4 GST-RhoA + DH + alkaline phosphatase

**Figure S6**. Pulldown experiments using obscurin $DH_{5681-5889}$ as a ligand and GST or GST-RhoA as a bait. Addition of EDTA or alkaline phosphatase lead to a nucleotide free state of RhoA.

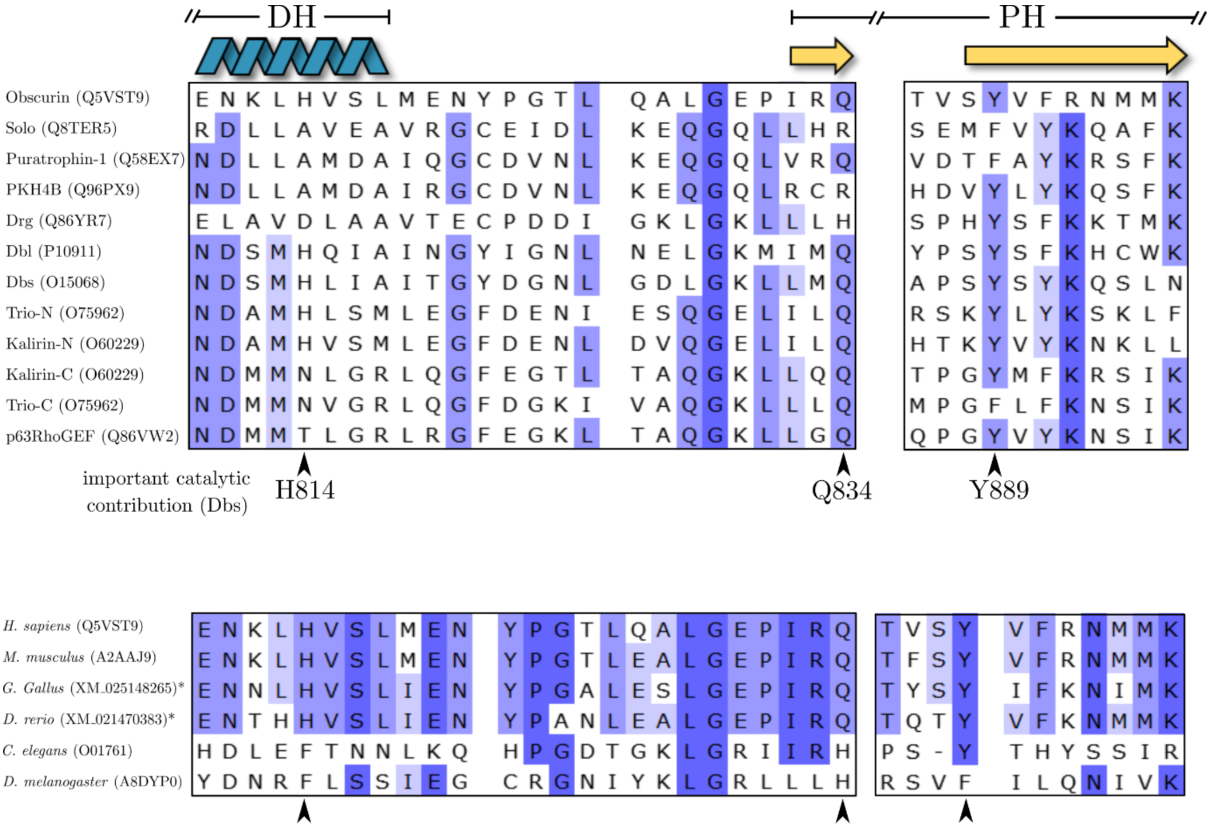


**Figure S7**. Sequence analysis of amino acids at the DH-PH domain interface. Black arrowheads indicate important catalytic contribution (numbering refers to Dbs sequence). Conserved residues are highlighted in blue with colorintensity highlighting the degree of conservation. **Top panel** shows comparison of human Trio-subfamily GEFs. **Bottom panel** shows obscurin DH-PH domain interface across different species. The unique Uniprot-identifier for each protein is given in brackets. When the protein was not available on Uniprot (*), the NCBI-Reference sequence is given instead.

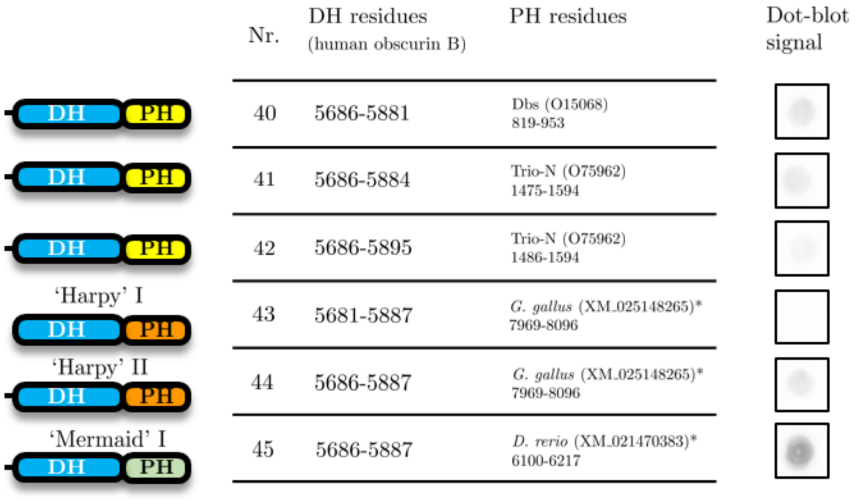


**Figure S8**. Chimeric obscurin RhoGEF fragments and dot-blot analysis of the eluted fraction after affinity purification of the indicated domains. The unique Uniprot-identifier for each protein is given in brackets. When the protein was not available on Uniprot (*), the NCBI-Reference sequence is given instead.

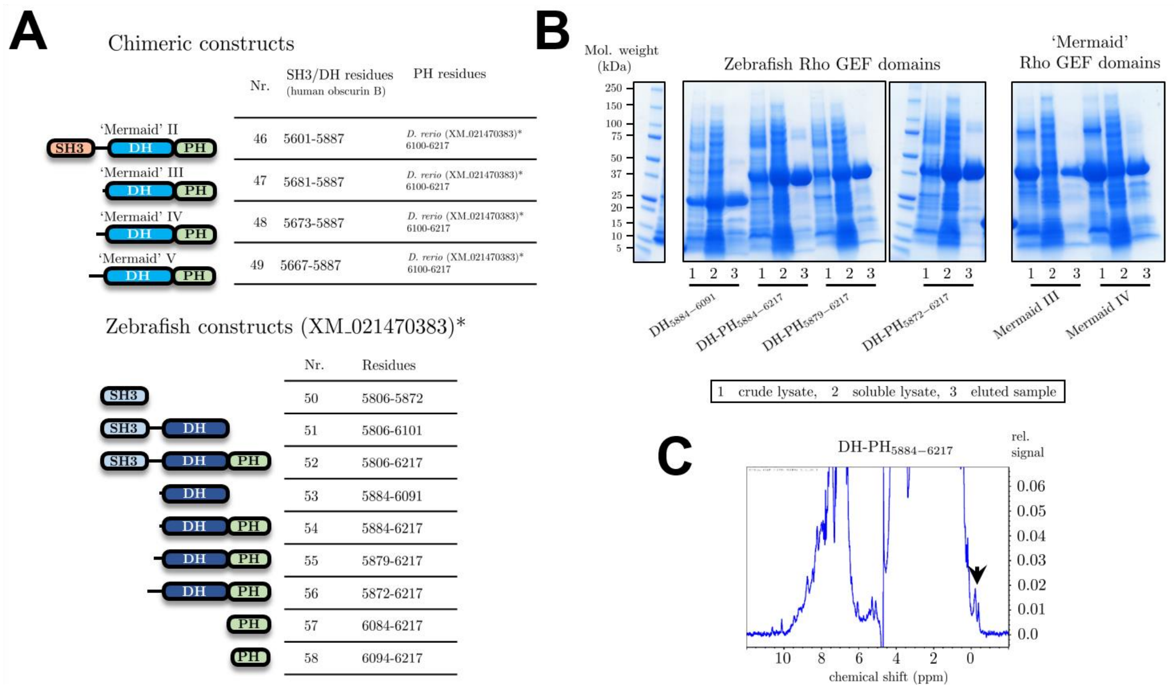


**Figure S9**. **A**, design of further human/zebrafish chimeras and zebrafish obscurin RhoGEF fragments. The unique Uniprot-identifier for each protein is given in brackets. When the protein was not available on Uniprot (*), the NCBI-Reference sequence is given instead. **B**, SDS-PAGE analysis after affinity purification of selected zebrafish and chimeric obscurin RhoGEF fragments. **C**, 1D-NMR analysis of purified zebrafish DH-PH$_{5884\text{-}6217}$ fragment. NMR spectrum exhibits wide peak dispersal and peaks below 0 ppm (black arrow), indicating that the protein is folded.

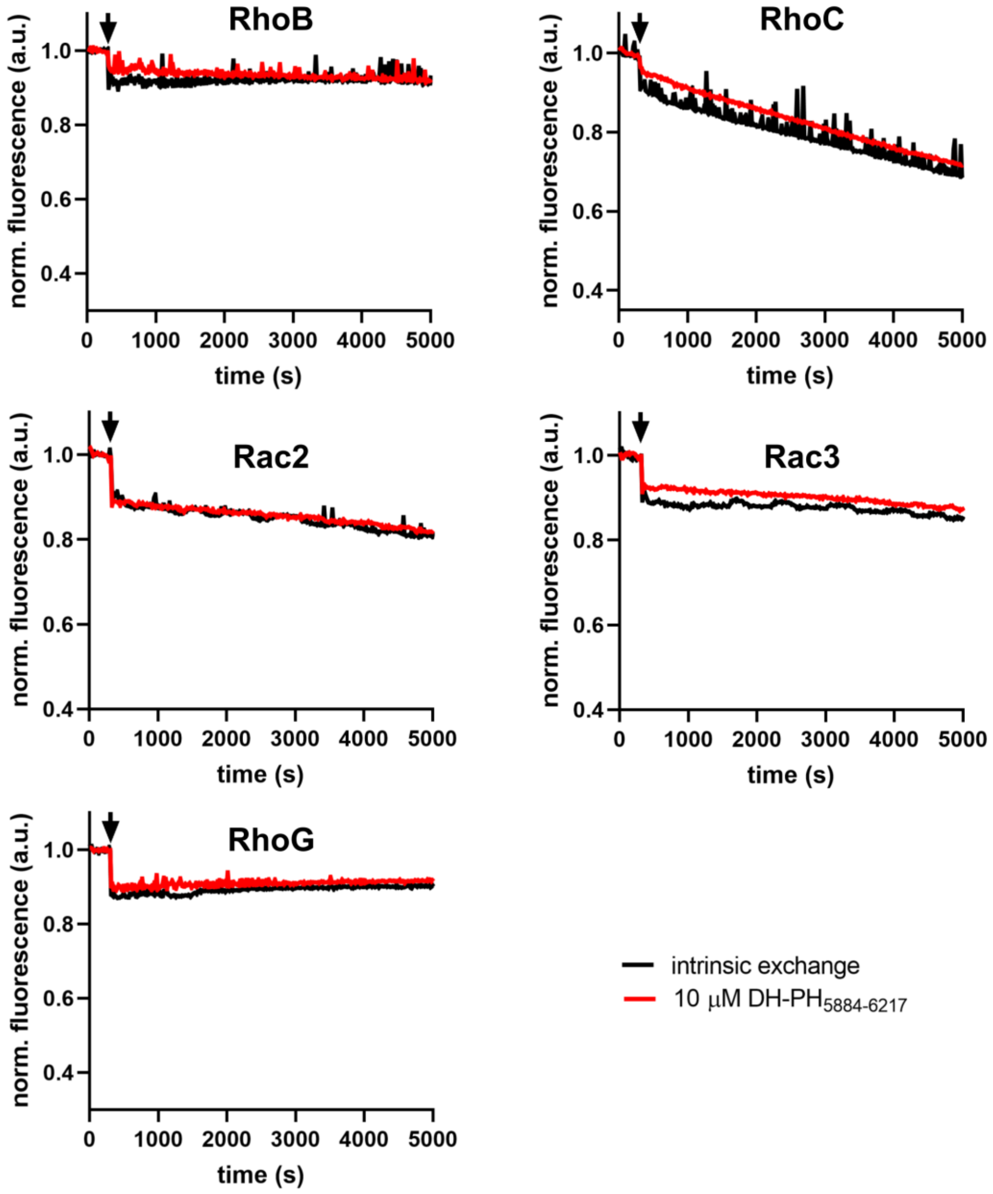


**Figure S10**. Guanosine nucleotide exchange factor activity of zebrafish obscurin DH-PH$_{5884\text{-}6217}$ towards RhoB, RhoC, Rac2, Rac3 and RhoG. Black arrows indicate addition of buffer/GEF/EDTA. Data represent mean of n = 2-3 experiments.

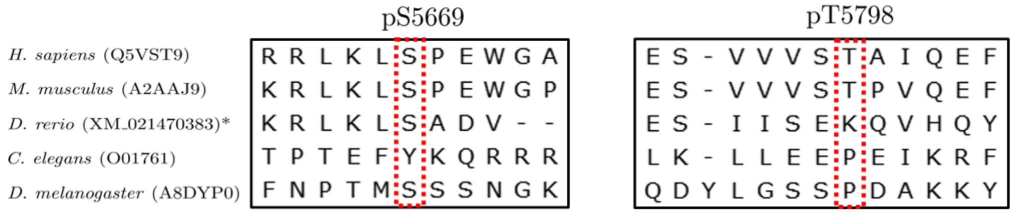


**Figure S11**. Conservation of phosphorylation sites in the human obscurin RhoGEF region reported by Potts et al. 2017. Numbering refers to human obscurin B sequence.

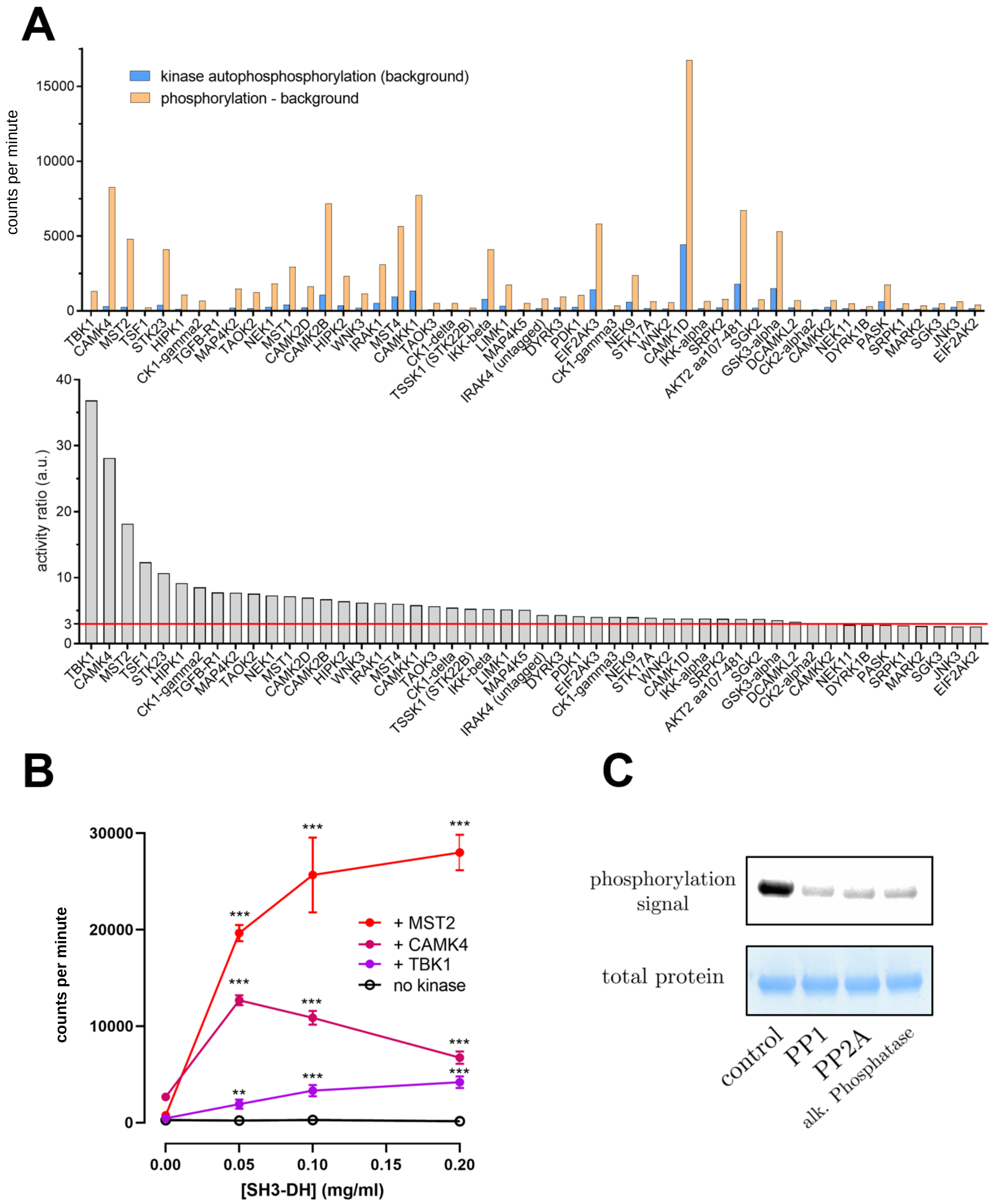


**Figure S12**. Top 50 results of a commercial kinase screen using [γ[33]]-ATP and human obscurin SH3-DH as substrates. **A**, (top panel) corrected absolute phosphorylation (orange) of the SH3-DH domain after exposure to a kinase next to the autophosphorylation background signal of that kinase (blue). (Bottom panel) top 50 hits of kinases sorted by the activity ratio of each experiment which considers the corrected absolute phosphorylation of the substrate relative to the phosphorylation background. A value >3 is considered a significant hit. **B**, validation experiment with same method of top 3 hits in screening assay at different substrate concentrations and n = 3 replicates per concentration (** $p < 0.01$, *** $p < 0.001$, students t-test vs no kinase condition). While MST2 addition resulted in strong and saturable phosphorylation, TBK1 led to much lower phosphorylation levels and CaMK4 addition led to an intermediate phosphorylation level exhibiting a biphasic behaviour with phosphorylation levels decreasing at higher substrate concentrations. Since MST2 showed the highest phosphorylation levels of obscurin SH3-DH, we focused on MST2 phosphorylation in all further experiments. **C**, ProQ[TM] diamond stain signal of MST2-phosphorylated obscurin SH3-DH after addition of no phosphatase (control), phosphatases PP1, PP2A or alkaline Phosphatase.

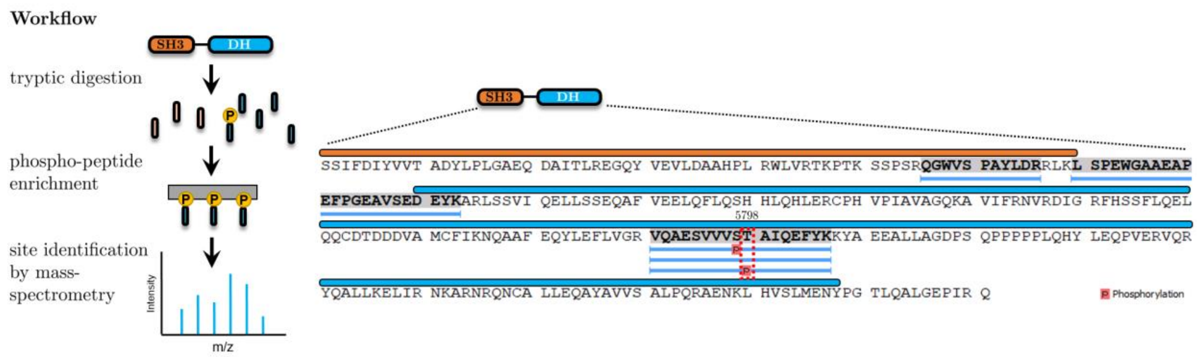


**Figure S13**. Identification of MST2 phosphorylation site within the human obscurin RhoGEF region via mass-spectrometric analysis of the digested phosphoprotein (workflow shown left). Protein sequence and identified peptides and phosphopeptides are shown on the right. Although both pSer5797 and pThr5798 peptides were identified, the precision of the identified site is often associated with an uncertainty of 1 or 2 residues. Since Thr5798 was observed to be phosphorylated *in vivo* (Potts et al., 2017), we concluded that the phosphorylation site is likely Thr5798.

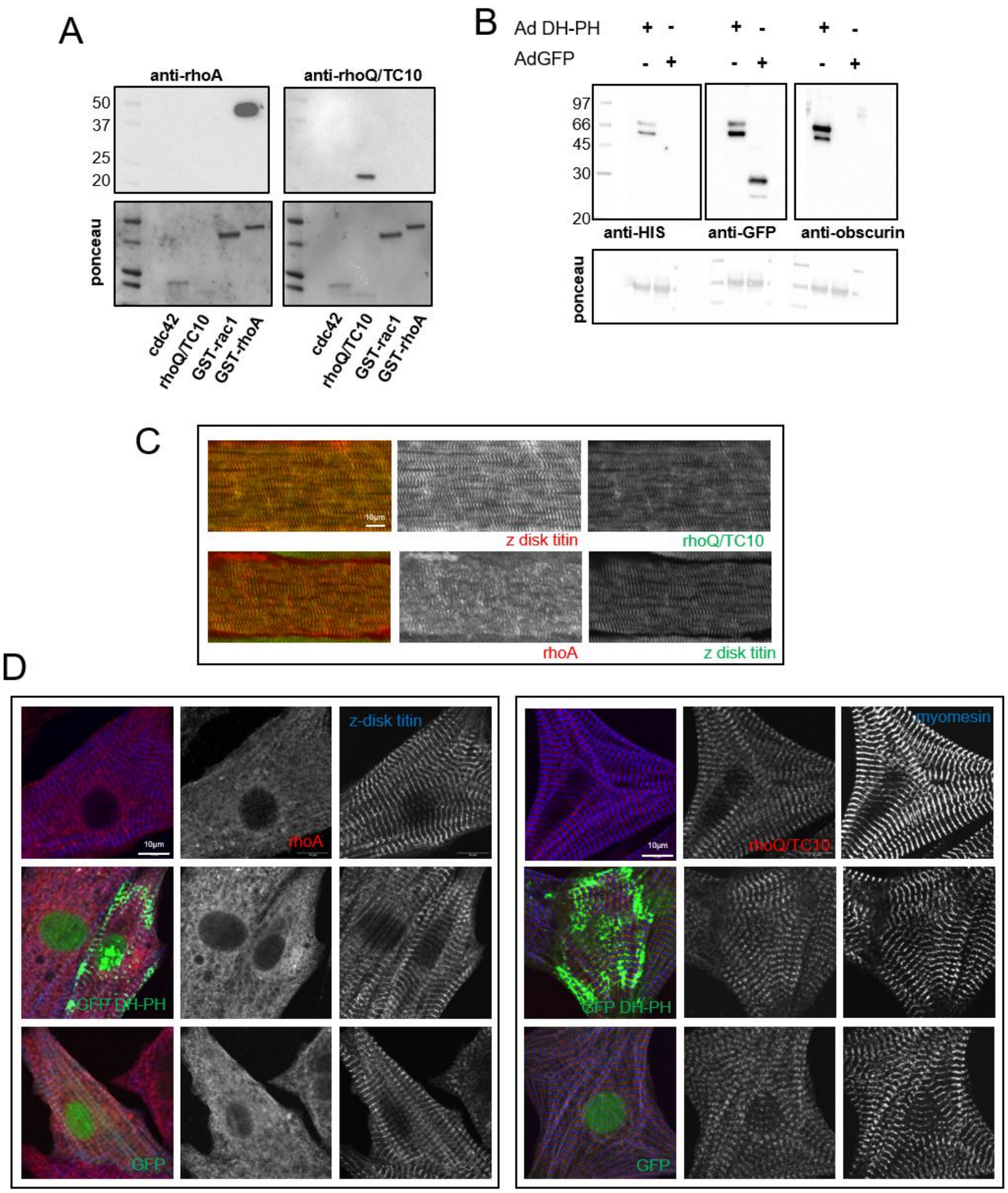


**Figure S14**. Validation experiments and localisation of RhoA and RhoQ/TC10. **A**, Western blot validation of RhoA and RhoQ/TC10 antibodies using recombinant GTPases shows that antibodies specifically bind their target epitope. **B**, **Upper**: Western blot confirming expression of obscurin DH-PH. HEK293 cells transduced with adenovirus containing GFP tagged obscurin DH-PH (Ad DHPH) or GFP alone (Ad GFP) and probed with anti-HIS, anti-GFP or anti-obscurin antibodies. **Lower**: ponceau stain. **C**, Mouse skeletal muscle (tibialis anterior) stained with **upper**: rhoQ/TC10 (green) and z-disk titin (red) or **lower**: rhoA (red) and z-disk titin (green). **D**, rhoA and rhoQ/TC10 localisation does not change upon overexpression of GFP tagged obscurin DH-PH in neonatal rat cardiomyocytes. **Upper row**: untransduced cells. **Middle row** (GFP DH-PH): cardiomyocytes transduced with adenovirus containing GFP obscurin DH-PH. **Bottom row** (GFP): cardiomyocytes transduced with adenovirus containing GFP only. **L**: rhoA (red), z-disk titin (blue). **R**: rhoQ/TC10 (red), myomesin (blue).

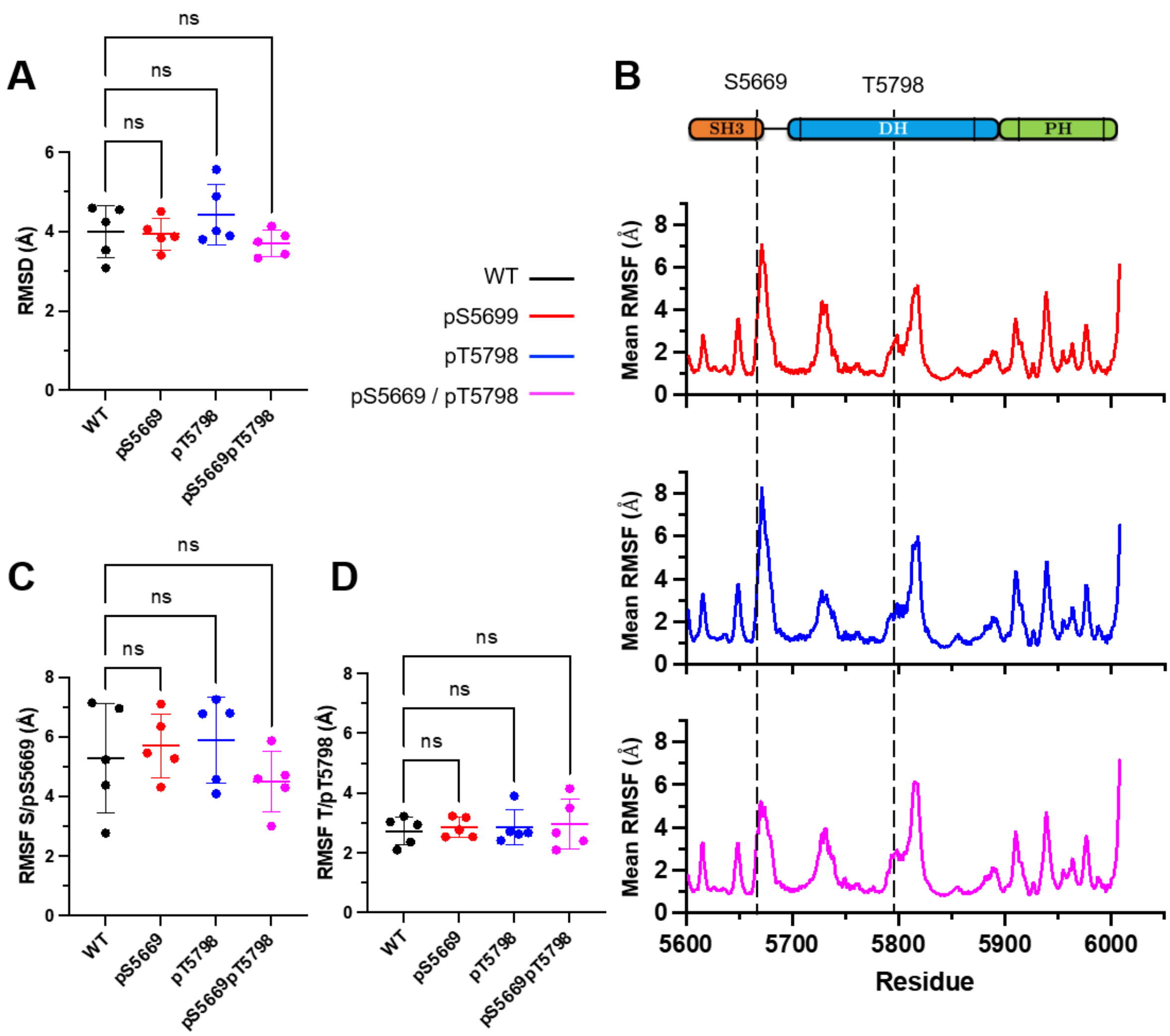


**Figure S15**. Molecular Dynamics simulations of Obscurin SH3-DH-PH WT and phosphorylated at either serine 5669, threonine 5798, or both. A total of n=5 runs was performed for each simulation. **A,** plot showing mean root-mean-squared deviation (RMSD) of structures compared to initial structure across simulation. The absence of significant differences in the RMSD between the four species suggest that phosphorylation at these residues does not induce large conformational changes or inter-domain motions. **B**, mean root-mean squared fluctuation (RMSF) per residue for SH3-DH-PH pS5669 (top), SH3-DH-PH pT5798 (middle) and SH3-DH-PH pS5699 / pT5798 (bottom), compared to WT. The position of the phosphorylated residues are indicated on the plots and indicate that the phosphorylated residues are in regions of high flexibility, in particular Ser5669. **C** and **D,** plots showing RMSF for residue 5669 (**C**) and 5798 (**D**) for the different molecular species. No significant differences between the molecular species were observed, including at the phosphorylated residues themselves. Similarly, we found no differences in RMSF for the conserved histidine (5879), glutamine (5899) and tyrosine (5945) residues (cf. Figure 3C in the main text) that would be expected to assist in the exchange reaction (data not shown).

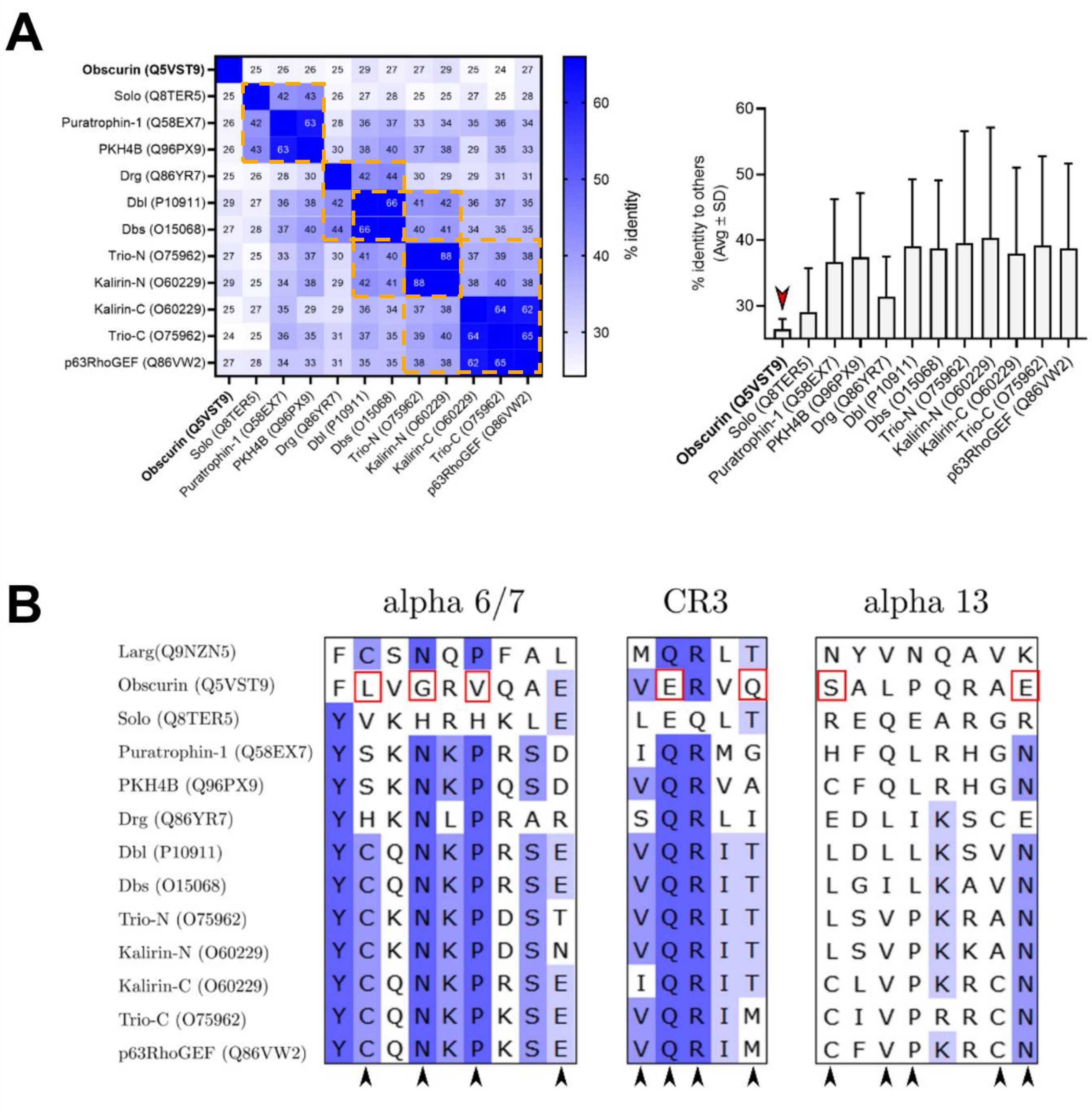


**Figure S16**. Bioinformatic analysis of the obscurin RhoGEF region amino acid sequence. **A**, multiple sequence alignments of the DH-PH domain sequences of all trio-subfamily members. Heatmap of the percent identity values on the left shows that obscurin is the only GEF that does not cluster (orange squares) with other subfamily members. We defined clusters as largest possible square neighborhoods along the diagonal that have at least 35% sequence identity. This is confirmed by plotting the average identity values for each GEF depicted in the right bar graph, showing that obscurin has the least average identity to other members of the trio-subfamily. **B**, analysis of key residues in the DH domain that have been identified to be constitutively involved in the interaction with GTPases from all Rho-subfamily members (Rho, Rac, Cdc42) in 13 DH/GTPase complex structures (Jaiswal et al., 2013). From 30 such functionally important and highly conserved residues (black arrowheads), obscurin shows significant differences at 7 of those residues (highlighted by red squares) in the regions alpha 6/7, CR3 and alpha 13 of the DH domain.
